# Redefining the role of Ca^2+^-permeable channels in photoreceptor degeneration using diltiazem

**DOI:** 10.1101/2020.12.04.411827

**Authors:** Soumyaparna Das, Valerie Popp, Michael Power, Kathrin Groeneveld, Christian Melle, Luke Rogerson, Marlly Achury, Frank Schwede, Torsten Strasser, Thomas Euler, François Paquet-Durand, Vasilica Nache

## Abstract

Hereditary degeneration of photoreceptors has been linked to over-activation of Ca^2+^-permeable channels, excessive Ca^2+^-influx, and downstream activation of Ca^2+^-dependent calpain-type proteases. Unfortunately, after more than 20 years of pertinent research, unequivocal evidence proving significant and reproducible photoreceptor protection with Ca^2+^-channel blockers is still lacking. Here, we show that both D- and L-cis enantiomers of the anti-hypertensive drug diltiazem were very effective at blocking photoreceptor Ca^2+^-influx, most probably by blocking the pore of Ca^2+^-permeable channels. Yet, unexpectedly, this block neither reduced the activity of calpain-type proteases, nor did it result in photoreceptor protection. Remarkably, application of the L-cis enantiomer of diltiazem even led to a strong increase in photoreceptor cell death. These findings shed doubt on the previously proposed links between Ca^2+^ and retinal degeneration and are highly relevant for future therapy development as they may serve to refocus research efforts towards alternative, Ca^2+^-independent degenerative mechanisms.

## I. INTRODUCTION

In the retina rod photoreceptors respond to dim light and enable night-time vision, whereas cone photoreceptors respond to bright daylight and enable colour vision. *Retinitis pigmentosa* (RP) is a group of hereditary diseases where rod primary degeneration is followed by secondary cone loss, ultimately leading to blindness (1, 2). *Achromatopsia* (ACHM) is a related disease where a genetic defect causes cone degeneration without significant rod loss (3). Regrettably, most cases of RP/ACHM remain without effective treatment, even though photoreceptor death has been linked to overactivation of Ca^2+^-permeable channels (4, 5).

Phototransduction in rods and cones intricately links Ca^2+^- and cGMP-signalling. cGMP levels are regulated by guanylyl cyclase, producing cGMP, and phosphodiesterase-6 (PDE6), hydrolysing cGMP. In darkness, cGMP opens the cyclic nucleotide-gated channel (CNGC), located in the photoreceptor outer segment (OS), causing influx of Ca^2+^ and Na^+^ (6). This influx is countered by the Na^+^-Ca^2+^-K^+^-exchanger (NCKX) in the OS and by the ATP-driven Na^+^-K^+^-exchanger (NKX) in the photoreceptor inner segment (IS) (6). As a result, the cell is depolarized at approximately −35 mV (7). The consequent activation of Cav1.4 (L-type) voltage-gated Ca^2+^-channels (VGCCs), located in the cell body and synapse, mediates further Ca^2+^ influx and synaptic glutamate release (7, 8). In light, PDE6 rapidly hydrolyses cGMP, leading to CNGC closure, Ca^2+^ decrease, and photoreceptor hyperpolarization. Subsequently, VGCC closes, ending synaptic neurotransmitter release.

Loss-of-function mutations in PDE6 lead to cGMP accumulation and CNGC overactivation, which may result in an abnormally strong influx of Ca^2+^ into photoreceptor OSs (9, 10) and sustained activation of VGCCs, mediating even more Ca^2+^ influx (11). In RP animal models, such as in the *Pde6b* mutant *rd1* and *rd10* mice (12), excessive Ca^2+^ is thought to lead to high activity of Ca^2+^-dependent calpain-type proteases and photoreceptor death (13, 14). In *rd1* animals the roles of CNGC and VGCC in photoreceptor cell death were studied by crossbreeding with knockouts (KO) of either CNGC (*Cngb1*^-/-^) or VGCC (*Cacna1f*^-/-^). While, VGCC KO did not influence *rd1* degeneration (15), CNGC KO strongly delayed *rd1* photoreceptor loss (16), highlighting CNGC as a target for pharmacological intervention.

Many studies over the past two decades have assessed the protective potential of Ca^2+^-channel blockers in photoreceptor degeneration (reviewed in (5)). The anti-hypertensive drug diltiazem is particularly interesting because its D-cis enantiomer blocks mostly VGCCs, while the L-cis enantiomer acts more strongly on CNGCs (17, 18). Both D- and L-cis-diltiazem have been suggested to delay *rd1* photoreceptor degeneration (11, 19, 20). However, other studies reported conflicting or contradictory results (21-23).

Here, we assessed the effect of D- and L-cis-diltiazem on heterologously expressed rod and cone CNGCs. We show that L-cis-diltiazem efficiently reduces rod CNGC activity in a voltage- and cGMP-dependent manner, most probably by obstructing its conductive pore. Surprisingly, in retinal cultures, derived from *rd1* and *rd10* mice, neither D-nor L-cis-diltiazem prevented photoreceptor degeneration. Rather, CNGC inhibition with L-cis-diltiazem exacerbated photoreceptor loss. Together, our results indicate that CNGC or VGCC inhibition effectively reduces photoreceptor Ca^2+^ levels, however, this will not decrease, but may instead increase, photoreceptor degeneration.

## II. RESULTS

### Differential effects of D- and L-cis-diltiazem on photoreceptor CNGC

To assess the effects of D- and L-cis-diltiazem on retinal CNGCs, we expressed the heterotetrameric rod CNGA1:B1a- and cone CNGA3:B3-channels in *Xenopus laevis* oocytes and examined their functional characteristics using electrophysiological recordings. We first confirmed correct assembly of heterotetrameric CNGC in the oocyte plasma membrane: (1) Co-expression of the main subunits, rod CNGA1 and cone CNGA3, with their modulatory subunits, CNGB1a and CNGB3, respectively, led to a strong increase of cAMP efficacy in heterotetrameric *vs*. homotetrameric channels (24, 25) (Fig. S1a,b). (2) Expression of CNGCs containing GFP-labelled CNGB1a or CNGB3 subunits and staining the oocyte membrane with fluorescently-labelled lectin (AlexaFluor™633-WGA) demonstrated plasma membrane localization of heterotetrameric channels (Fig. S1b,e).

We measured next the CNGC concentration-activation relationships in the presence of cGMP (Table S1; Fig. S1c,f). Under physiological conditions, at −35 mV and with up to 5 μM cGMP (26), CNGC activity reached ~6 % of its maximum for cones and ~1 % for rods (Fig. 1a-d). When applied to the intracellular side of the membrane, neither D-nor L-cis-diltiazem (up to 100 μM) significantly influenced physiological CNGC activity (Fig. 1, Table S2). In the presence of saturating cGMP (3 mM), both diltiazem enantiomers inhibited cone and rod CNGCs (grey areas in Fig. 1a-d). The strongest effect on both CNGC isoforms was triggered by L-cis-diltiazem, while rod CNGC was most sensitive to both D- and L-cis-diltiazem (Fig. 1e, Table S2). With pathologically high cGMP (100 μM), emulating RP-like conditions, the diltiazem effect on rod CNGCs, mirrored closely our observations in the presence of 3 mM cGMP (Fig. 1e,f).

**Figure 1:**
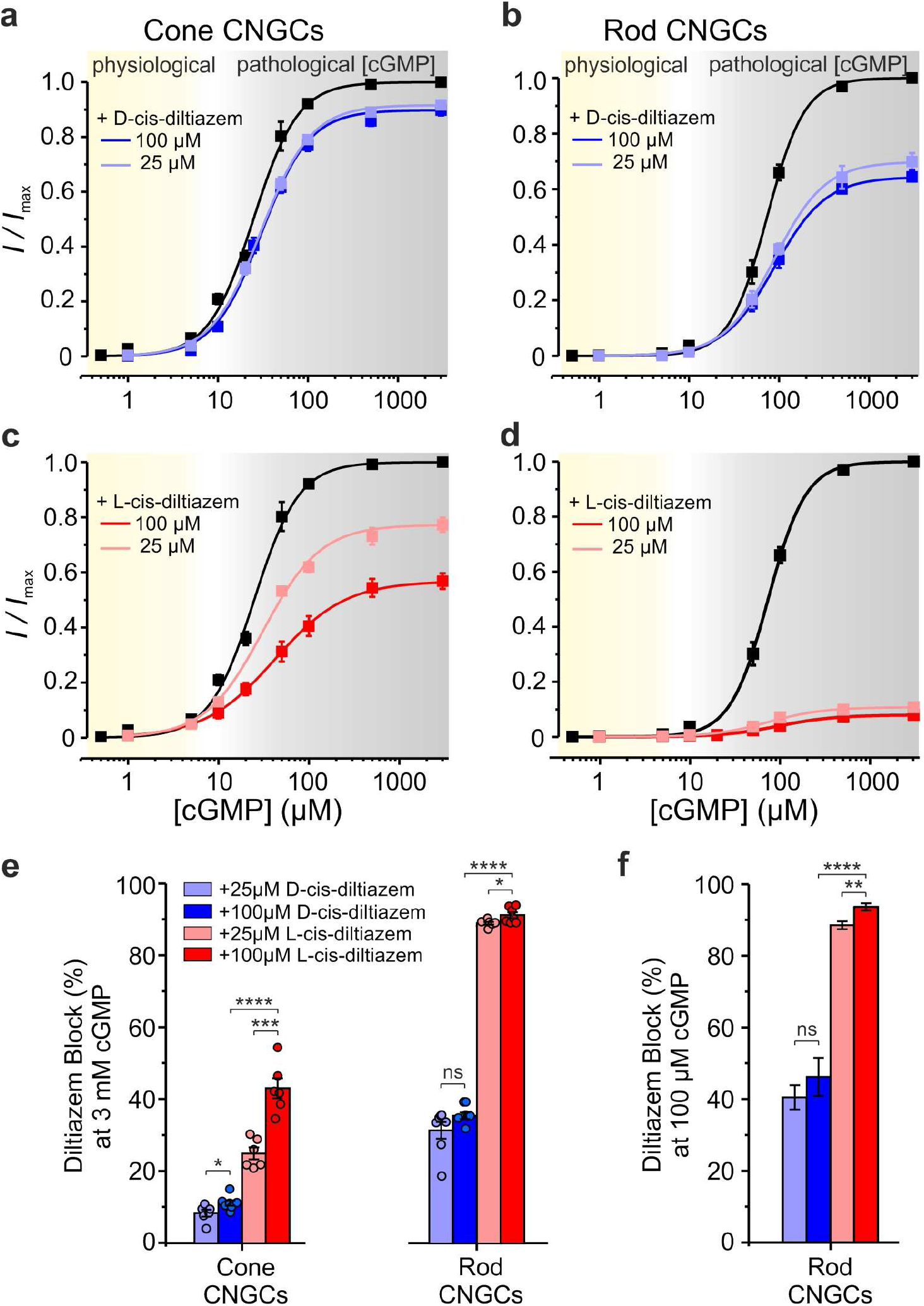
Effects of D- and L-cis-diltiazem on rod and cone CNGC activity. (**a-d**) Concentration-activation relationships for heterotetrameric cone (**a, c**) and rod (**b, d**) CNGCs in the presence of either 100 μM or 25 μM D- or L-cis-diltiazem, respectively, measured at −35 mV. The respective curves represent fits of the experimental data points with the Hill equation (Eq. 1). Black symbols show the normalized cGMP-triggered current amplitudes in the absence of diltiazem and are shown to point out the effect of the blocker. Light- and dark-blue symbols represent data obtained in the presence of D-cis-diltiazem, at 25 and 100 μM, respectively (**a, b**). Light- and dark-red symbols represent data obtained in the presence of L-cis-diltiazem at 25 and 100 μM, respectively (**c, d**). (**e, f**) D- and L-cis-diltiazem - block (%, ±SEM) of CNGCs in the presence of 3 mM (**e**) and 100 μM cGMP (**f**), respectively. The amount of diltiazem block was calculated using Eq. 2 (see Materials and Methods). The respective symbols represent single measurements (see also Table S1 and S2).

Both diltiazem enantiomers (at 100 μM) showed a stronger inhibitory effect at depolarizing (+100 mV) than at hyperpolarizing (−100mV) membrane voltages: for D-cis-diltiazem by a factor of ~4 and ~7, and for L-cis-diltiazem by a factor of ~2.6 and ~1.4 in case of cone and rod CNGC, respectively (Fig. S2 and S3). In addition, we observed a voltage-dependent increase of the *EC*_50_-values with a maximum at +100 mV and a systematic decrease of the *H*-values at all tested voltages (Table S1 and S3, Fig. S2 and S3c,d). D- and L-cis-diltiazem showed similar effects on *EC*_50_- and *H-* values, suggesting that both diltiazem enantiomers reduced the CNGC apparent affinity and the cooperativity between their subunits through a similar mechanism.

In conclusion, (1) under physiological conditions, neither D-nor L-cis-diltiazem affect CNGC activity; (2) at saturating cGMP-concentration, diltiazem had a differential voltage-dependent effect, with a stronger inhibition of rod- *vs*. cone-CNGCs, the effect of L-exceeding that of D-cis-diltiazem, and with maximal inhibition at depolarizing voltages.

### Influence of D- and L-cis-diltiazem on CNGC gating kinetics

We then studied the influence of diltiazem on CNGC gating kinetics (Fig. 2a,b). When applying cGMP and diltiazem simultaneously, the inhibition occurred only after channel activation, suggesting that diltiazem blocked open channels only. When cGMP and diltiazem were simultaneously removed, the channel deactivation was considerably delayed, indicating that diltiazem hindered channel closure.

**Figure 2:**
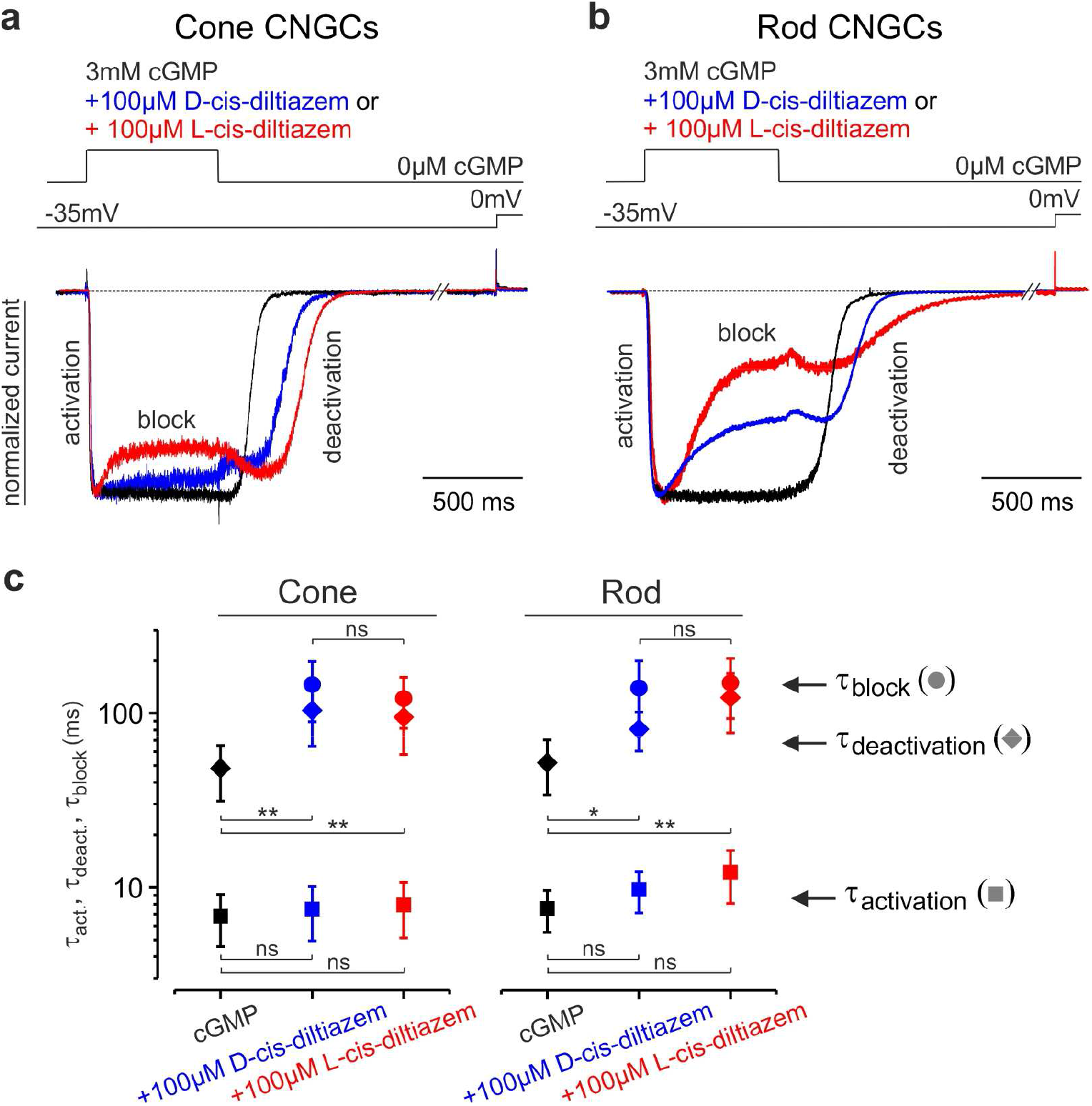
D- and L-cis-diltiazem influence rod and cone CNGC gating kinetics. Superimposition of representative activation-, deactivation- and block-time courses following a concentration jump from 0 μM cGMP to either 3 mM cGMP or 3 mM cGMP + 100 μM D- or L-cis-diltiazem and back to 0 μM cGMP for cone (**a**) and rod (**b**) CNGCs (n=5-9). The current traces (blue for D-, red for L-cis-diltiazem) were normalized to the initial current level triggered by 3 mM cGMP (black) in the absence of diltiazem. Above the current traces are depicted the experimental protocols. The small current increase observed during washout onset mirrors the initial phase of the diltiazem removal. **c**) CNGC-activation, -deactivation and -block time constants (τ_act_, τ_deact_, τ_block_). The respective traces in **a**) and **b**) were fitted with mono-exponential functions (Eq. 3) and the resulting mean time constants and statistical analysis (ms, ±SEM) were included in Table S4. The time course of channel deactivation was fitted starting after the initial delay due to diltiazem removal.

The activation time course of rod and cone CNGCs (τ_act_) seemed unaffected by diltiazem, whereas the channel’s deactivation (τ_deact_) was delayed and slowed down by a factor of ~2 (Fig. 2c, Table S4). Also, the kinetics of the blocking event was similar for both channel isoforms (τ_block_). This suggested a common blocking mechanism for D- and L-cis-diltiazem, possibly by obstructing the CNGC pore. We next tested whether the observed diltiazem-induced block was Ca^2+^-dependent (27, 28). In the presence of extracellular Ca^2+^ (1 mM CaCl_2_) we found a reduced cGMP-triggered activation of CNGCs (Fig. S4a), an effect that was consistent with a very slow Ca^2+^ permeation (29). Nevertheless, the influence of Ca^2+^ on the strength of the L-cis-diltiazem-induced block was only minor (Fig. S4b), indicating that Ca^2+^ did not prevent diltiazem binding to its binding pocket.

### Influence of L-cis-diltiazem on cGMP binding

To assess whether diltiazem influences ligand binding, we employed confocal patch-clamp fluorometry (cPCF) (30, 31). Here, we used rod CNGC, the most diltiazem-sensitive channel, L-cis-diltiazem, the enantiomer with the strongest blocking effect and f*cGMP (8-[DY-547]-AHT-cGMP), a fluorescent derivative of cGMP (Fig. 3) (32). As expected, the f*cGMP-induced current (10 μM) was reduced in the presence of L-cis-diltiazem to 10.8 ± 1.1%. Upon blocker removal from an open channel, the recovery of CNGC activity showed two steps with different kinetics: a fast and a very slow phase which took several minutes (Fig. 3). Surprisingly, this behaviour differed from the faster diltiazem washout observed when the blocker and cGMP were concomitantly removed (Fig. 2a,b). This indicated an acceleration of diltiazem unbinding triggered by simultaneous channel closure.

**Figure 3:**
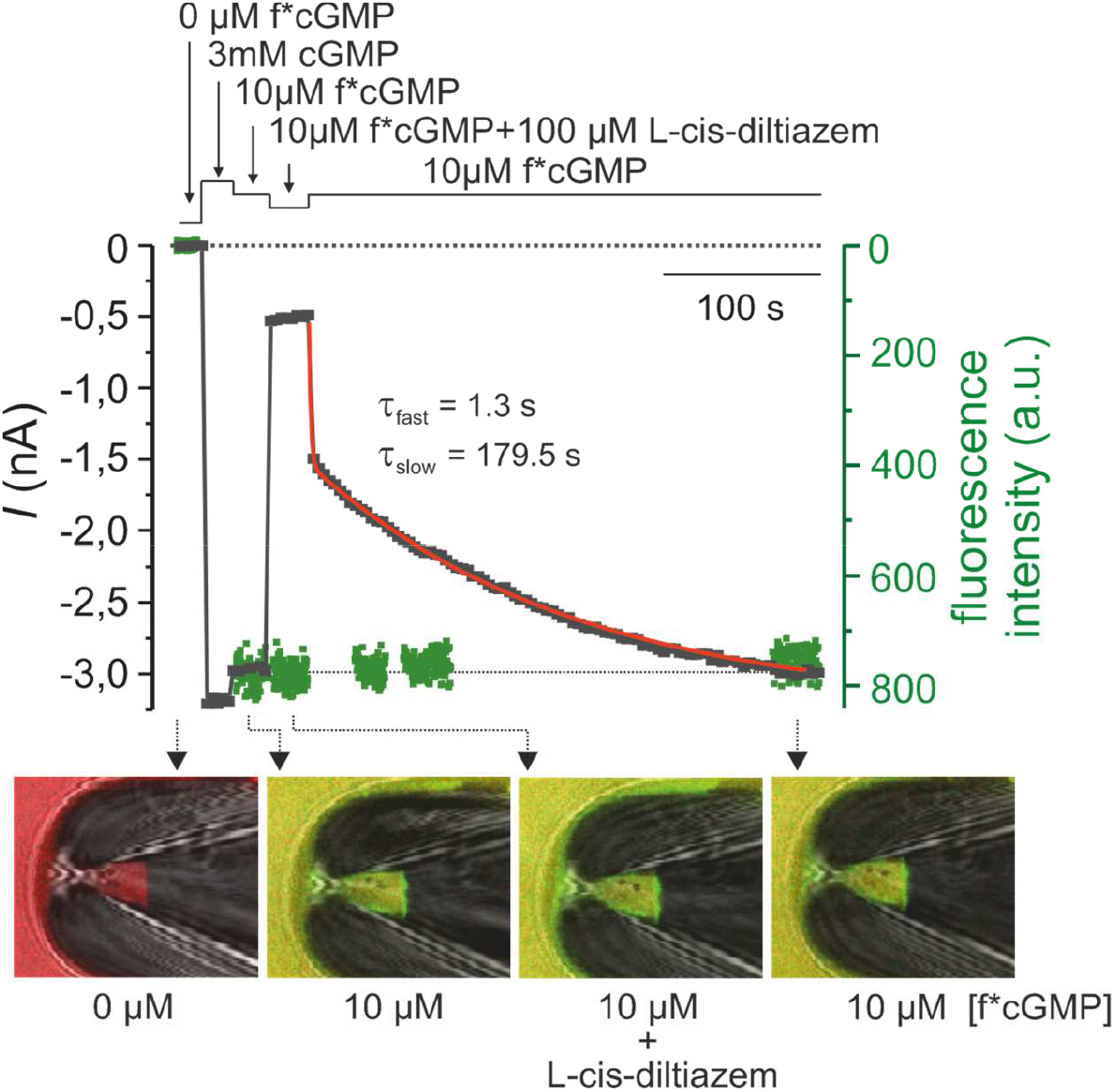
L-cis-diltiazem does not influence cGMP binding to rod CNGCs. Shown is a representative cPCF measurement for studying simultaneously f*cGMP (8-[DY-547]-AHT-cGMP) binding and rod CNGCs activation in the presence of 100 μM L-cis-diltiazem. f*cGMP has a higher potency than cGMP: 10 μM f*cGMP triggered already 87.4 ± 1.4% activation of rod CNGC, which is ~20 times more than the activation triggered by 10 μM cGMP. The experimental protocol is depicted above the diagram. Black symbols represent the current amplitude measured under steady-state conditions. Green symbols represent the f*cGMP fluorescence signal which indicates the amount of ligand binding. The steady-state binding signal was normalized to the level of the 10 μM f*cGMP-induced current. The lower part of the diagram shows confocal images of glass pipettes, containing CNGCs-expressing membrane patches, which were obtained during the measurement in the absence (first image, left), in the presence of 10 μM f*cGMP (second and fourth image) and in the presence of 10 μM f*cGMP + 100 μM L-cis-diltiazem (third image). The time course of the current recovery upon removal of L-cis-diltiazem was fitted with a double exponential function yielding τ_fast_ = 1.5 ± 0.1 s and τ_slow_ = 161.9 ± 24.5 s (red line, n=8, Eq. 4).

During the application of L-cis-diltiazem and after its removal, we observed no major change in the intensity of the fluorescence signal which encodes for the total amount of bound f*cGMP to CNGCs (Fig. 3). This showed that L-cis-diltiazem inhibits CNGCs independent of cGMP binding, in line with our electrophysiological data on the channel’s apparent affinity (Fig. S3c,d).

### Effects of D- and L-cis-diltiazem on light induced photoreceptor Ca^2+^ responses

We next recorded light-induced photoreceptor Ca^2+^-responses using two-photon imaging and transgenic mice expressing a fluorescent Ca^2+^-biosensor exclusively in cones (13). As the biosensor was absent from OS, we recorded from cone terminals (Fig. 4a), using synaptic Ca^2+^ signals as a proxy for changes in membrane potential caused by light-dependent OS CNGC modulation (14). We presented series of 1-s flashes of light and measured the change (decrease) in terminal Ca^2+^, quantifying the responses using area-under-the-curve (AUC), without (control) and with diltiazem enantiomers at different concentrations (25, 50, 100 μM).

**Figure 4:**
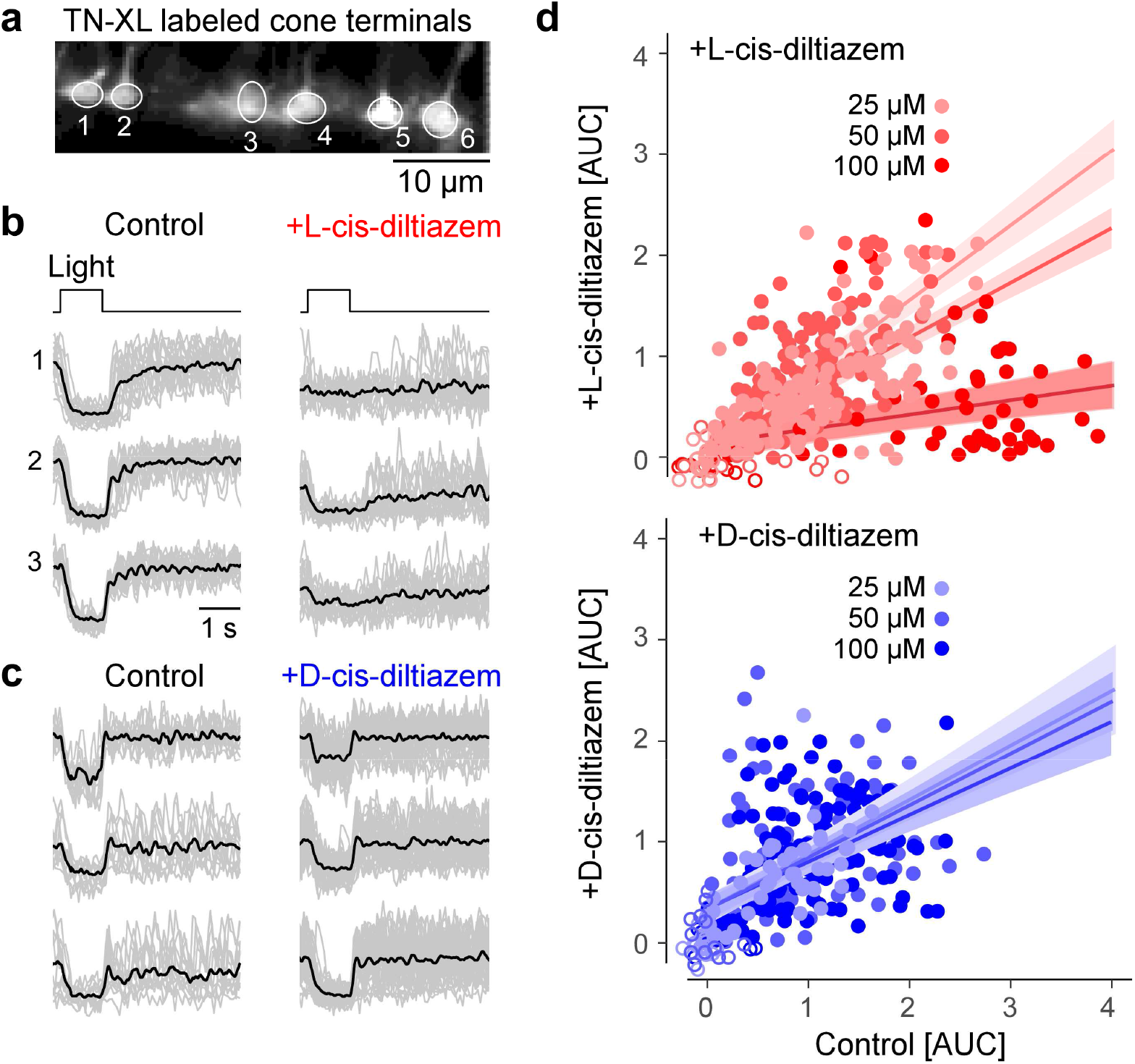
Light-evoked Ca^2+^-responses are reduced by L- but not by D-cis-diltiazem. (**a**) Recording of light-evoked Ca^2+^-responses from cone photoreceptor terminals, in mouse retinal slices expressing the fluorescent Ca^2+^ sensor TN-XL in cones. (**b, c**) Exemplary Ca^2+^ responses before (control) and in the presence of 100 μM L- (**b**) or D-cis-diltiazem (**c**) (grey, single trials; black, mean of n trials, with control in (b), n=13; control in (c), n=19; L-cis-diltiazem, n=19; D-cis-diltiazem, n=38). (**d**) Scatter plot of response size (as area-under-the-curve, AUC) for both D-cis (blue; 25/50/100 μM n=137/138/61 cells) and L-cis-diltiazem (red; 25/50/100 μM n=62/140/162 cells; each data point represents a cell). Fits show mean predictions and standard errors from a multivariate linear model (Table S5).

We used a multivariate linear model to identify what factors (*i.e*., enantiomer, concentration) were significant for predicting cellular responses (Tables S5). This analysis revealed that L-cis-diltiazem significantly decreased responses in a concentration-dependent manner, whereas D-cis-diltiazem did not affect light-induced cone Ca^2+^ responses (Fig. 4b-d; for detailed statistics, see Table S5). L-cis-diltiazem (but not D-cis-diltiazem) also tended to decrease the Ca^2+^-baseline level (Fig. 4b,c; left *vs*. right).

These data suggested that, at physiological cGMP concentrations, treatment with L-cis-diltiazem locked synaptic Ca^2+^ concentrations at a low level, abolishing cone light responses. D-cis-diltiazem, on the other hand, had no significant effect on light-induced Ca^2+^ responses in cones. Since in heterologously expressed CNGCs (Fig. 1) the rod channel isoform was more sensitive to L-cis-diltiazem than its cone counterpart, L-cis-diltiazem likely reduces rod Ca^2+^ levels even more.

### Expression of CNGCs in photoreceptor outer segments

The effects of Ca^2+^-channel inhibitors on photoreceptor viability were tested using wild-type (wt) and *rd1* mice. To ascertain that CNGC was expressed in *rd1* retina in the relevant timeframe, we performed immunostaining for the CNGB1a channel subunit on retinal tissue sections collected at six different time-points between post-natal day (P) 11 and 30 (Fig. S5a-c). The CNGB1a staining was also used to estimate OS length (Fig. S5d). In wt retina OS length quadrupled between P11 and P30, while dramatically decreasing in *rd1* retina in the same time window. As a proxy for photoreceptor degeneration, we measured the thickness of the outer nuclear layer (ONL), which contains the photoreceptor cell bodies, within a timeframe that includes most of the unfolding of *rd1* photoreceptor degeneration (P11 to 30). Linear mixed effect models revealed statistically significant effects of genotype and post-natal day, showing significant differences for OS length between wt and *rd1* with increasing age. (Table S6). At the beginning of the *rd1* degeneration (~P10), OS length as assessed by CNGB1a expression was still comparable to that in wt animals (average P11 OS length least square means difference between wt and *rd1*: 0.18 ± 2.05 μm, *F* (1, 25.25) = 0.0078; *p* = 0.9304). Hence, in *rd1* retina the window-of-opportunity for CNGC-targeting treatments was expected to last until P11 at least.

### Proteolytic activity in photoreceptors after treatment with D- and L-cis-diltiazem

The influx of Ca^2+^ through CNGCs may be connected to activation of Ca^2+^-dependent calpain-type proteases (33). Therefore, we investigated the effects of D- and L-cis-diltiazem treatment, using an *in situ* calpain activity assay and immunodetection of activated calpain-2 (34), and organotypic retinal explant cultures derived from wt and *rd1* animals, treated from P7 to P11.

In wt retina, least square means plot showed calpain activity and calpain-2 activation to be rather low, when compared to *rd1* where both markers labelled large numbers of photoreceptors in the ONL (Fig. 5a-d). In both genotypes, treatment with D-cis-diltiazem had no detectable effect on the numbers of photoreceptors positive for calpain activity or calpain-2 activation (Fig. 5e-h, Tables S7, S8). Surprisingly, when retinal explants were treated with L-cis-diltiazem (Fig. 5i-l), calpain-2 activation in the ONL was significantly increased (*F* (1, 11.87) = 14.7372; *p* = 0.0024; Fig. 5i,j,n; Table S8). Thus, neither D-nor L-cis-diltiazem reduced overall calpain activity in wt or *rd1* photoreceptors, while a significant increase in calpain-2 activation was observed with L-cis-diltiazem (Fig. 5m,n; Tables S7, S8).

**Figure 5:**
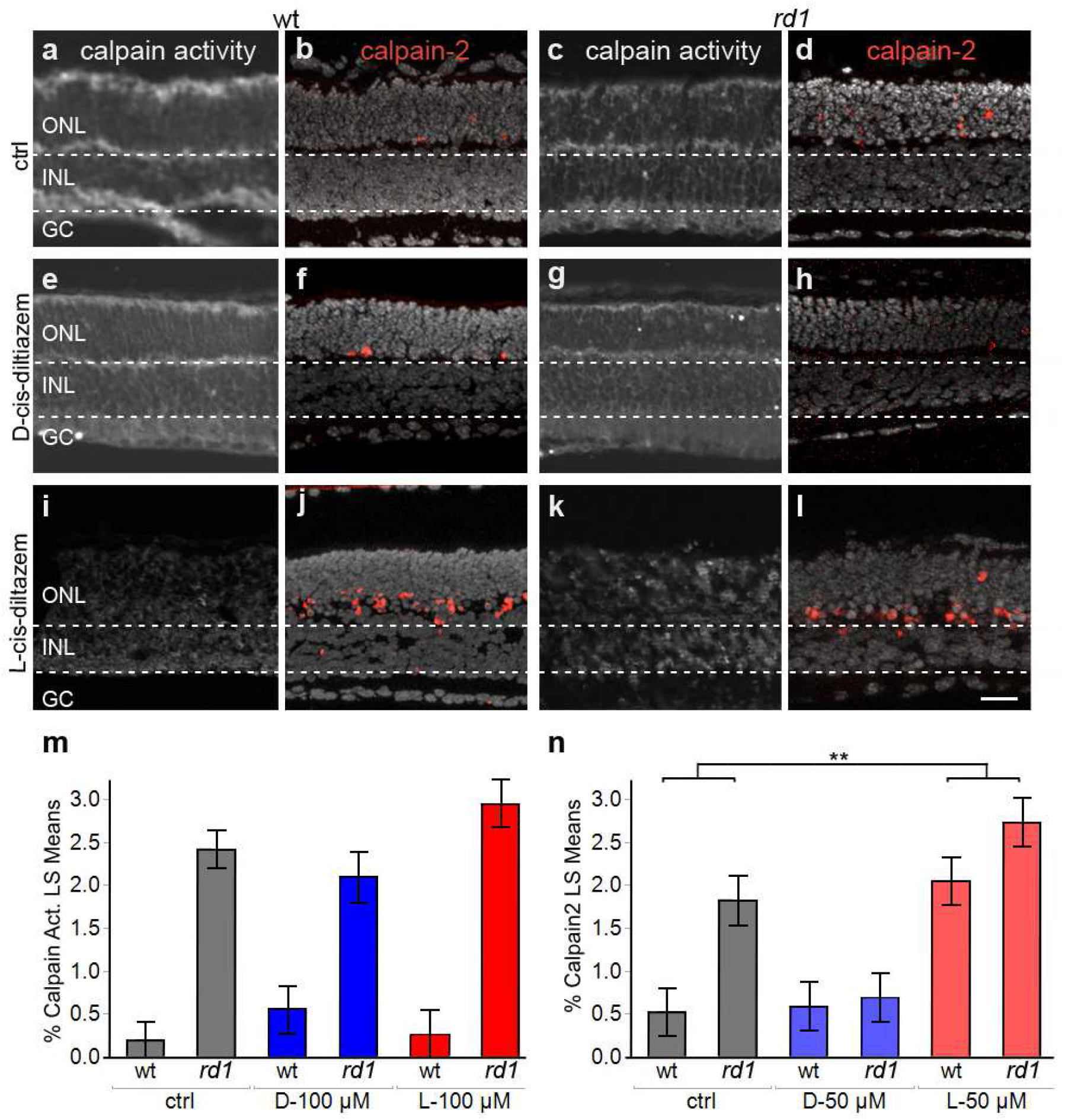
Effects of diltiazem treatment on calpain activity. Calpain-activity assay and immunostaining for activated calpain-2 in wt and *rd1* retina. Untreated retina (ctrl; **a-d**) was compared to treatment with D-cis diltiazem (**e-h**) or L-cis diltiazem (**i-l**). The bar graphs show the least-square (LS) means percentages of cells positive for calpain activity (**m**) and activated calpain-2 (**n**) in wt and *rd1* retina, compared to the untreated control (ctrl). Asterisks indicate a statistically significant difference from a contrast test performed between control and 50 μM L-cis-diltiazem treatment (L-50 μM). For statistical analysis, see Tables S7 and S8; error bars represent SEM; ** = *p* < 0.01. Scale bar =30 μm.

Activity of caspase-type proteases is commonly associated with apoptosis. To investigate possible links with apoptosis, *rd1* treated and untreated retinas were tested for activation of caspase-3, using an antibody directed against the active protease (35). Under all conditions tested, caspase-3 activity was essentially undetectable in retinal sections (Fig. S6, Table S7), thus ruling out an important contribution of caspase activity and, by extension, of apoptosis to retinal degeneration, with or without diltiazem treatment.

### Impact of D- and L-cis-diltiazem on *rd1* photoreceptor degeneration

We used the TUNEL assay to quantify photoreceptor cell death (36) after D- or L-cis-diltiazem treatment on organotypic retinal explant cultures derived from wt, *rd1*, and *rd10* animals (33). The ONL of wt retinal explants displayed a relatively low number of TUNEL positive cells, when compared to *rd1* (Fig. 6a,b). D-cis-diltiazem treatment did not elevate the numbers of dying cells in wt or *rd1* ONL (Fig. 6c,d). In contrast, L-cis-diltiazem (Fig. 6e,f) increased cell death in both wt (*F* (1, 21.63) = 86.7207, *p* < 0.0001) and *rd1* retina (*F* (1, 26.68) = 191.1994, *p* < 0.0001; Fig. 6g,h, Tables S7, S8).

**Figure 6:**
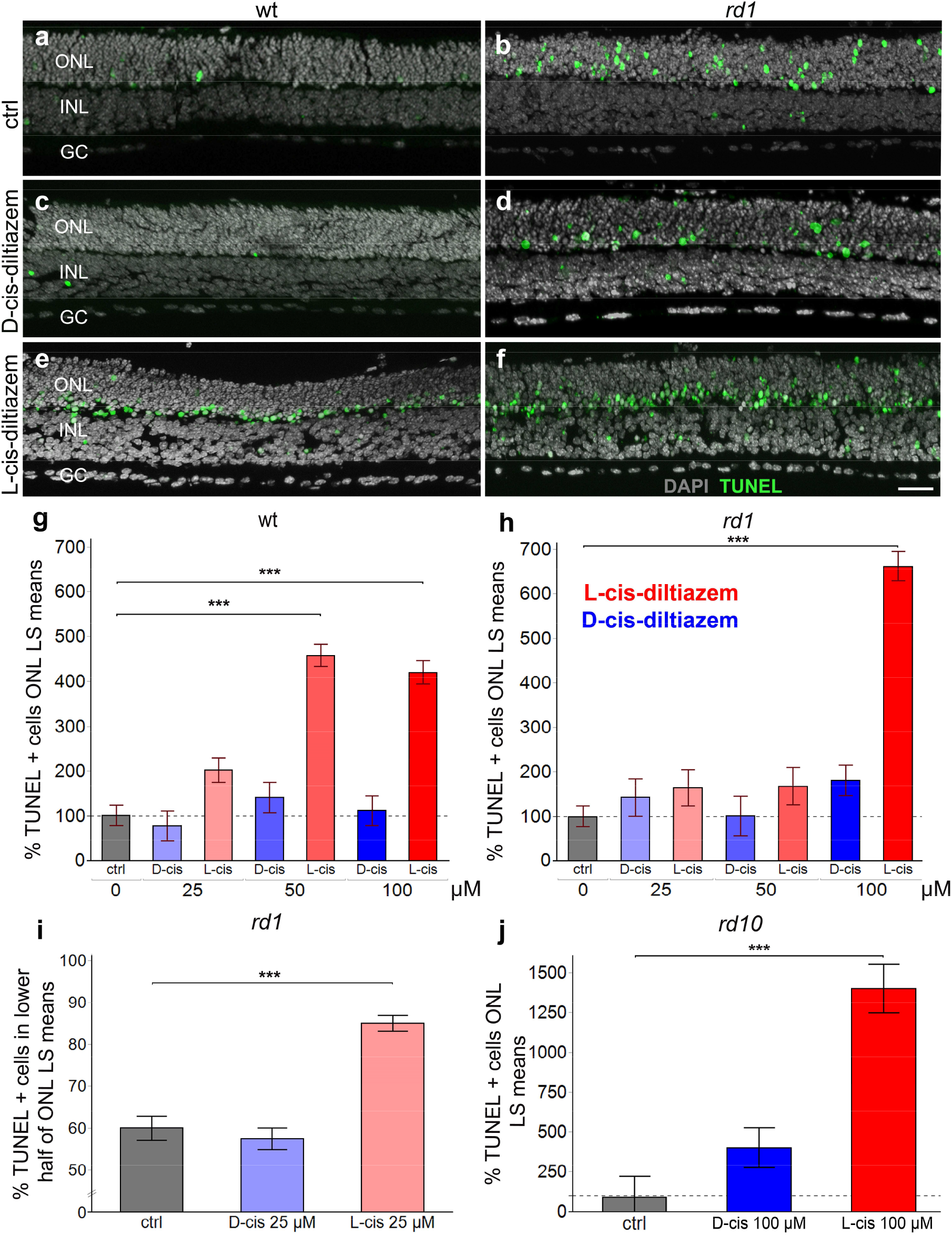
Effects of D- and L-cis-diltiazem on retinal cell viability. The TUNEL assay was used to label dying cells (green) in wt and *rd1* retinal explant cultures. DAPI (grey) was used as a nuclear counterstain. Control retina (untreated; **a, b**) was compared to retina treated with either 50 μM of D- (**c, d**) or L-cis-diltiazem (**e, f**). Note the large numbers of dying cells in the *rd1* outer nuclear layer (ONL). The bar charts show the least-square (LS) means percentage of TUNEL positive cells as a function of diltiazem concentration, for wt (**g**) and *rd1* (**h**) retina, as a function of localization with respect to the outer plexiform layer (OPL) (**i**), and for *rd10* retina (**j**), respectively. In wt, *rd1*, and *rd10* retina treatment with L-cis-diltiazem strongly increased the numbers of TUNEL positive cells in the ONL. Statistical significance was analysed by post-hoc contrast test (*cf*. Table S8), errors bars represent SEM, *** = *p* < 0.001. INL = inner nuclear layer, GC = ganglion cell layer. Scale bar = 50μm.

Typically, TUNEL positive cells in untreated retinal explants were uniformly distributed across the whole ONL (Fig. 6a,b; percent *rd1* dying cells in inner half of ONL = 60 ± 3%). With D-cis-diltiazem treatment 57 ± 4% of dying cells were localized to the same space (Fig. 6c,d), while, curiously, with L-cis-diltiazem treatment 85 ± 3% (*F* (1, 10.42) = 54.2025, *p* < 0.0001; Tables S7, S8) of the TUNEL positive cells were seen in the lower ONL half (Fig. 6e,f,i).

A study on *rd10* retina yielded results similar to *rd1*: 100 μM D-cis-diltiazem had no significant effect on cell death, while 100 μM L-cis-diltiazem treatment caused a strong increase in photoreceptor cell death (*F* (1, 9.25) = 42.9966, *p* < 0.0001; Fig. 6j).

Taken together, our data indicate that L-cis-diltiazem treatment in wt retina was toxic to photoreceptors at concentrations above 25 μM. In *rd1* retina showed such toxicity only at concentrations above 50 μM, which might be related to higher photoreceptor cGMP levels and concomitant higher CNGC activity. In comparison, D-cis-diltiazem, up to 100 μM, did not detectably influence cell viability in either wt-, *rd1*-, or *rd10-* retinas. Importantly, both diltiazem enantiomers failed to show protective effects in *rd1* or *rd10* mutant retina.

## III. DISCUSSION

We show that diltiazem enantiomers were highly effective at blocking photoreceptor Ca^2+^ influx through CNGCs at pathologically high cGMP concentrations, likely by blocking the channel’s pore. Unexpectedly, this block did not result in photoreceptor protection. These results raise the question whether Ca^2+^-permeable channels are suitable targets for therapeutic interventions, and furthermore suggest that high intracellular Ca^2+^ is not *per se* a driver of photoreceptor death.

### Effect of D- and L-cis-diltiazem on photoreceptor CNGCs

At physiological cGMP, neither D- nor L-cis-diltiazem showed an appreciable inhibitory effect on heterologously expressed CNGCs. At high, RP-like cGMP concentrations, both diltiazem enantiomers reduced rod and cone CNGC activity, although L-cis-diltiazem had a much stronger inhibitory effect on rod CNGC than D-cis-diltiazem. The inhibition was strongly voltage dependent, suggesting that a disease-induced photoreceptor depolarization would amplify diltiazem effects on CNGCs. Although electrophysiological recordings from single photoreceptors of retinal disease models are rare (37), *rd1* rod photoreceptors can be expected to be permanently depolarized due to elevated CNGC activity triggered by high cGMP.

Earlier studies on photoreceptor CNGC proposed several binding sites for diltiazem, either at the pore entrance, on the cytoplasmic side of the channel (38), or within the channel pore (39). Recently, L-cis-diltiazem was shown to bind within the conductive pathway of voltage-gated Ca^2+^-channels (Ca_v_Ab and Ca_v_1.1) (40, 41). Our data on CNGC, *e.g*., (1) the time delay observed between channel activation and diltiazem block and between diltiazem removal and channel closure, (2) the acceleration of diltiazem removal by a concomitant channel deactivation, and (3) the undisturbed cGMP binding in the presence of diltiazem, suggests that L-cis-diltiazem acts in a similar way, by blocking the CNGC pore. These findings, indicating an open-channel block, concur with the recent eukaryotic (42) and human CNG (43) cryo-EM channel structures.

Moreover, we observed a negative influence of diltiazem on the cooperativity between CNGC subunits. Since L-cis-diltiazem inhibits only heterotetrameric channels (25), this suggests a direct interaction between diltiazem and the modulatory subunits, rod CNGB1a and cone CNGB3, respectively. Future studies using molecular-docking approaches may help identify the diltiazem binding site within the channel’s pore and its biophysical characteristics.

### An overview on Ca^2+^ flux in photoreceptor degeneration

Photoreceptor degeneration in hereditary retinal diseases has long been proposed to be caused by excessive Ca^2+^ influx (10, 19), *i.e*. the “high Ca^2+^ hypothesis”. Paradoxically, too low Ca^2+^ was also suggested to cause photoreceptor death, something that may be called the “low Ca^2+^ hypothesis” (44). Subsequently, we discuss Ca^2+^ flux in different photoreceptor compartments (Fig. 7a) and will attempt to resolve some of the contradictions between high and low Ca^2+^ hypotheses.

**Figure 7:**
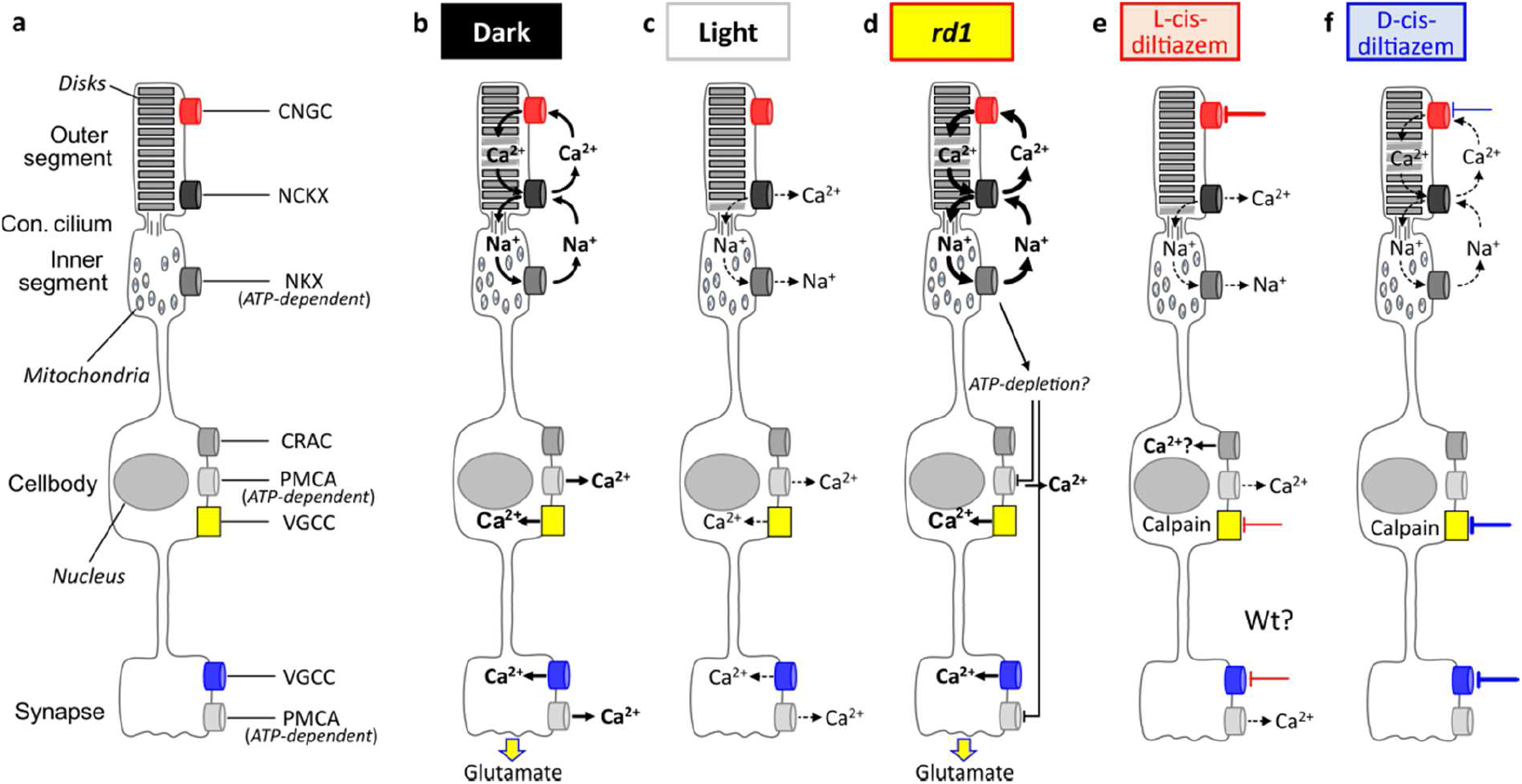
Schematic representation of photoreceptor Ca^2+^ flux under different experimental conditions. (**a**) The phototransduction cascade is compartmentalized to the photoreceptor outer segments, which harbour cyclic nucleotide-gated channel (CNGC) and Na^+^/Ca^2+^/K^+^ exchanger (NCKX). The connecting cilium links outer to inner segment, which holds almost all mitochondria and the ATP-driven Na^+^/K^+^ exchanger (NKX). The cell body harbours the nucleus as well as Ca^2+^-release activated channel (CRAC), plasma membrane Ca^2+^-ATPase (PMCA), and voltage-gated Ca^2+^ channels VGCC. PMCA and VGCC are also found in the synapse. (**b**) In the dark, the flux of Na^+^ and Ca^2+^ ions across the photoreceptor membrane (*i.e*., the dark current) keeps the cell in a continuously depolarized state. The Ca^2+^ ions enter the outer segment via CNGC and exits via NCKX. The Na^+^ gradient needed to drive NCKX is maintained by the ATP-dependent NKX in the inner segment. At the same time, in the photoreceptor cell body and synapse, VGCC allows for Ca^2+^ influx, mediating synaptic glutamate release. In the cell body and synapse, Ca^2+^ is extruded by the ATP-dependent PMCA. (**c**) In light, CNGC closes, while Ca^2+^ continues to exit the cell via NCKX, leading to photoreceptor hyperpolarization. This in turn closes VGCC, ending synaptic glutamate release. (**d**) In *rd1* photoreceptors, high cGMP continuously opens CNGC, representing a situation of “constant darkness”. Excessive NKX activity in *rd1* may cause a depletion of ATP, preventing Ca^2+^ extrusion via PMCA. (**e**) L-cis-diltiazem (red lines) blocks predominantly CNGC, with an additional block on VGCC at high concentrations. This resembles a situation of “constant light” and may cause a depletion of intracellular Ca^2+^ and secondary Ca^2+^ influx via activation of CRAC. (**f**) D-cis-diltiazem (blue lines) inhibits predominantly VGCC, with a partial block on CNGC at high concentrations.

In wt photoreceptors, under dark conditions, Ca^2+^ and Na^+^ enter the OS (Fig. 7b). While Ca^2+^ is extruded from the OS via NCKX, Na^+^ is actively exported by ATP-dependent NKX in the IS (45). In cell body and synapse, Ca^2+^ is extruded by the plasma membrane Ca^2+^-ATPase (PMCA) (46). During illumination, cGMP levels drop, CNGCs and then VGCCs close, and Ca^2+^ levels in OS and synapse decrease (Fig. 7c). In *rd1* retina, the constitutively open CNGCs allow for permanent Ca^2+^ and Na^+^ influx, increasing NKX activity and perhaps resulting in ATP depletion (Fig. 7d) (47). L-cis diltiazem will block CNGC, reducing Ca^2+^ influx into the OS (Fig. 7e). D-cis-diltiazem in turn blocks mostly VGCC and prevents Ca^2+^ influx into cell body and synapse (Fig. 7f).

Surprisingly, calpain-2 activation was increased by the L-cis-diltiazem treatment. This may have been caused by a depletion of Ca^2+^ in intracellular stores and a subsequent activation of store-operated Ca^2+^ entry (SOCE) via Ca^2+^ release-activated Ca^2+^ channels (CRACs) (48). Indeed, VGCC block with diltiazem was recently shown to activate SOCE in vascular smooth muscle cells (49) and this process may selectively activate calpain-2 (50). Thus, photoreceptor degeneration initially caused by very low Ca^2+^ levels, may trigger a consequent increase of Ca^2+^ and calpain-2 activity via SOCE, possibly explaining the apparent contradiction between high and low Ca^2+^ hypotheses. Moreover, low photoreceptor Ca^2+^ levels will disinhibit guanylyl cyclase, increasing cGMP production (51), which may then kill photoreceptors independent of Ca^2+^ via over-activation of cGMP-dependent protein kinase G (PKG) (52, 53).

### Calpain activity, Ca^2+^, and cell death

Previously, we had proposed calpain activation in dying photoreceptors to be mediated by high cGMP-activated CNGCs (33). However, the low mobility of Ca^2+^ ions from OS to IS (13, 54, 55) argues against CNGC-dependent Ca^2+^-influx directly activating calpain in the cell body and beyond. Instead, our data suggests that calpain activation may be mediated indirectly by Ca^2+^ influx via VGCC located in cell body and synapse, in line with data obtained from genetic inactivation of VGCC (15).

L-cis-diltiazem was highly effective at blocking Ca^2+^ influx in heterologously-expressed rod CNGC and displayed a good rod *vs*. cone CNGC selectivity. When *rd1* retina was treated with L-cis-diltiazem, rod CNGCs were expressed in photoreceptors during the treatment period. Yet, we were unable to demonstrate a protective effect of L-cis-diltiazem on *rd1* retina. Even worse, at higher concentrations, L-cis-diltiazem showed obvious signs of toxicity, in wt, *rd1*, and *rd10* retina. This is contradicting the results seen with genetic inactivation of CNGC in *rd1* * *Cngb1*^-/-^ double-mutant mice (9). Yet, such animals likely still retain CNGA1 homotetrameric channels, which may allow for limited Ca^2+^ influx into the photoreceptor (56). In addition, loss-of-function mutations in CNGC genes are known to cause photoreceptor degeneration in both RP (57) and ACHM (58). Hence, on a genetic level, low activity of CNGC and decreased Ca^2+^ influx into photoreceptor OSs is clearly connected to photoreceptor degeneration.

Incidentally, inhibition of VGCC with D-cis-diltiazem also failed to show significant photoreceptor protection. This is in line with a number of earlier studies (reviewed in (5)) and corroborates on a pharmacological level our previous study employing the *rd1* * *Cacna1f*^-/-^ double-mutants, *i.e. rd1* mice in which the synaptic VGCC was dysfunctional (15). In contrast to L-cis-diltiazem, D-cis-diltiazem did not appear to be overly retinotoxic even at high concentrations. This corresponds to the genetic situation where loss-of-function mutations in VGCC impair synaptic transmission from photoreceptors to second order neurons. While such mutations can cause night-blindness, they do not usually cause photoreceptor degeneration (59).

Excessive Ca^2+^ influx via CNGC and/or VGCC has for a long time been suggested as a major driver for photoreceptor cell death (9, 19). However, follow-up studies have produced contradictory results (5). Our present study sheds light onto this enigma and demonstrates that both D- and L-cis enantiomers of the anti-hypertensive drug diltiazem can reduce photoreceptor Ca^2+^ influx. Remarkably, treatment with either compound and inhibition of either VGCC or CNGC did not result in photoreceptor protection. Moreover, the use of L-cis-diltiazem and the concomitant reduction of Ca^2+^ influx strongly reduced photoreceptor viability, indicating that Ca^2+^-influx was in fact protective, rather than destructive. Altogether, this supports the “low Ca^2+^” hypothesis (44) and cGMP-dependent processes (53) as the more likely causes of photoreceptor degeneration.

## IV. Materials and Methods

### Animals

Animals used in this study were handled according to the German law on animal protection. All efforts were made to keep the number of animals used and their suffering to a minimum. Mice were bred in the Tübingen Institute for Ophthalmic Research specified-pathogen-free (SPF) housing facility, under 12h/12h light/dark cycle, had *ad libitum* access to food and water, and were used irrespective of gender. The experimental procedures involving animals were reviewed and approved by the institutional animal welfare committee of the University of Tübingen.

For retinal explant cultures C3H/HeA *Pde6b ^rd1/rd1^* animals (*rd1*) and their respective congenic wild-type C3H/HeA *Pde6b* ^+/+^ counterparts (*wt*) were used (60). Further studies were performed on explants derived from C57BL/6J *Pde6b ^rd10/rd10^* animals (*rd10*) (33). For studying light-induced Ca^2+^ responses in cone photoreceptors, we used transgenic mice expressing the Ca^2+^ biosensor TN-XL (61) under the human red opsin promoter HR2.1 on a C57BL/6J background (13).

The procedures regarding the *Xenopus laevis* frogs and the handling of the oocytes had approval from the authorized animal ethics committee of the Friedrich Schiller University Jena (Germany). The respective protocols were performed in accordance with the approved guidelines.

### Molecular biology and functional expression of heterotetrameric CNGCs in *Xenopus laevis* oocytes

The coding sequences for the retinal CNGC subunits, bovine CNGA1 (NM_174278.2) (62) and CNGB1a (NM_181019.2) (63) from rod photoreceptors and human CNGA3 (NM_001298.2) (64) and CNGB3 (NM_019098.4) (65) from cone photoreceptors, were subcloned into the pGEMHE vector (66) for heterologous expression in *Xenopus laevis* oocytes. The surgical removal of oocytes was performed from adult frog females under anaesthesia (0.3% tricaine; MS-222, Pharmaq Ltd., Fordingbridge, UK). The oocytes were treated with collagenase A (3 mg/ml; Roche Diagnostics, Mannheim, Germany) for 105 min in Barth’s solution containing (in mM) 82.5 NaCl, 2 KCl, 1 MgCl_2_, and 5 HEPES, pH 7.5. After this procedure, oocytes of stages IV and V were manually dissected and injected with the genetic material encoding for the CNGC from rod and cone photoreceptors. For efficient generation of heterotetrameric channels, the ratio of CNGA3 mRNA to CNGB3 mRNA was 1:2.5 (25) and of CNGA1 mRNA to CNGB1a mRNA was 1:4 (67). After injection, the oocytes were kept at 18°C for 2 to 7 days in Barth’s solution containing (in mM) 84 NaCl, 1 KCl, 2.4 NaHCO_3_, 0.82 MgSO_4_, 0.41 CaCl_2_, 0.33 Ca(NO_3_)_2_, 7.5 Tris, cefuroxime (4.0 μg×ml^-1^), and penicillin/streptomycin (100 μg×ml^-1^), pH 7.4.

### Electrophysiology

Macroscopic ionic currents were measured with the patch-clamp technique and the inside-out configuration, using an Axopatch 200B patch-clamp amplifier (Axon Instruments, Foster City, CA). Recordings were made at room temperature. Current data were acquired using PATCHMASTER software (HEKA Elektronik, Lambrecht, Germany) with a sampling frequency of 5 kHz, and low-pass filtered at 2 kHz. From a holding potential of 0 mV, currents were elicited by voltage steps to −65 mV, then to −35 mV, and back to 0 mV. When mentioned, also voltage steps to −100 mV and +100 mV were recorded. The patch pipettes were pulled from borosilicate glass tubing (outer diameter 2.0 mm, inner diameter 1.0 mm; Hilgenberg GmbH, Germany). The initial resistance was 0.6-1.3 MΩ. Intracellular and extracellular solutions contained 140 mM NaCl, 5 mM KCl, 1 mM EGTA, and 10 mM HEPES (pH 7.4). The Ca^2+^-containing solutions were: 120 mM NaCl, 3 mM KCl, 2 mM NTA, 0.5 mM niflumic acid, 10 mM HEPES and 1 mM CaCl_2_ (pH 7.4) for the extracellular side and 145 mM KCl, 8 mM NaCl, 2 mM NTA, 10 mM HEPES and 0.05 mM CaCl_2_ (pH 7.4) for the intracellular side (28).

The cyclic nucleotides, cAMP (Merck KGaA, Darmstadt, Germany) or cGMP (Biolog LSI GmbH & Co KG, Bremen, Germany), were added to intracellular solutions as indicated. Either D- or L-cis-diltiazem (Abcam - ab120260 and Abcam - ab120532, respectively, Germany) were added to the cGMP-containing solutions to a final concentration of 25 μM and 100 μM as required. The diltiazem-containing solutions were prepared from stock solutions (10 mM) immediately before experiments. The cGMP-solutions, or solution mixtures containing cGMP and either D- or L-cis-diltiazem were administered via a multi-barrel application system to the cytosolic face of the patch.

For studying CNGC activation and deactivation kinetics we performed fast solution jumps (from zero to either 3 mM cGMP or 3 mM cGMP + 100 μM D- or L-cis-diltiazem and back to zero) by means of a double-barrelled θ-glass pipette mounted on a piezo-driven device (68). The recording rate was 20 Hz. The solution exchange at the pipette tip was completed within 1 ms (69).

### Confocal patch-clamp fluorometry (cPCF)

The influence of D- and L-cis-diltiazem on cGMP binding was studied by means of cPCF. The method has been described in detail previously (31, 70, 71). The experiments were performed in inside-out macropatches of *Xenopus laevis* oocytes expressing heterotetrameric rod CNGC, at −35 mV. As fluorescent ligand we used 8- [DY-547]-AHT-cGMP (f*cGMP). f*cGMP was prepared in analogy to the related cyclic nucleotides 8-[DY-547]-AET-cGMP and 8-[DY-547]-AHT-cAMP (31, 32). To be able to differentiate between the fluorescence of the bound f*cGMP from the fluorescence generated by the free f*cGMP in the bath solution, we used an additional red dye, DY647 (Dyomics, Jena, Germany), at a concentration of 1 μM.

Recordings were performed with an LSM 710 confocal microscope (Carl Zeiss Jena GmbH, Germany) and were triggered by the ISO3 hard- and software (MFK, Niedernhausen, Germany; sampling rate 5 kHz, 4-pole Bessel filter set to 2 kHz). Due to the relative long duration of the experiment, to avoid cell-membrane exposure to damaging amounts of light, binding was measured under steady-state conditions, during pre-selected time windows only: in the presence of 10 μM f*cGMP, during the jump to 10 μM f*cGMP + 100 μM L-cis-diltiazem and during L-cis-diltiazem removal from the open channels.

### Colocalization experiments

To verify the correct incorporation of heterotetrameric CNGCs into the oocyte plasma membrane we labelled the cone CNGB3- and rod CNGB1a-subunit by fusing enhanced GFP to their intracellularly located C terminus. At first, we introduced an *Avr*II site in pGEMHE-CNGB1a by site-directed mutagenesis at CNGB1a K1205 which was thereby changed to R1205. Afterwards, the PCR amplified *EGFP* gene was ligated into the newly generated *AvrII* site of the pGEHME-CNGB1a construct. To fuse EGFP into CNGB3 C-terminus, we introduced an *Xho*I site in pGEMHE-CNGB3 by site-directed mutagenesis at CNGB3 P668 and K669 which were thereby changed to L668 or E669, respectively. Afterwards, the PCR amplified *EGFP* gene was ligated into the newly generated *Xho*I site of the pGEHME-CNGB3 construct. The correct insertion of PCR products was confirmed by DNA sequencing.

The oocyte membrane was stained from the extracellular side with fluorescently labelled lectin (Alexa Fluor™ 633 - wheat germ agglutinin (Alexa-WGA), Invitrogen Life Technologies Corporation, Eugene, Oregon, red fluorescence signal) (71). For this the oocytes were incubated in 5 μg/ml Alexa-WGA for 7 minutes. Alexa-WGA was excited with the 633-nm line of a helium neon laser. GFP was excited with the 488-nm line of an argon laser. GFP- and WGA-fluorescence profiles measured along an imaginary line perpendicular to the plasma membrane, were quantified using the LSM 710 image analysis software.

### Analysis of the oocyte data

For concentration-activations relationships, each patch was first exposed to a solution containing no cGMP and then to a solution containing the saturating concentration of 3 mM cGMP. After subtracting the baseline current from the current amplitude in the presence of cGMP, the current response for each ligand concentration was normalized to the saturating current. The experimental data points were fitted using the Hill equation:

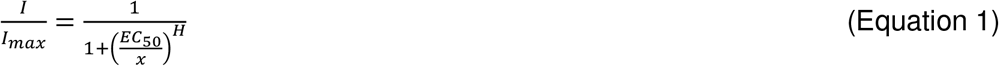

where *I* is the current amplitude, *I*_max_ is the maximum current induced by a saturating cGMP concentration, *x* is the ligand concentration, *EC*_50_ is the ligand concentration of half maximum effect, and *H* is the Hill coefficient. The analysis was performed with OriginPro 2016G software (OriginLab Corporation, Northampton, USA). Experimental data are given as mean ±SEM.

The effect of either D- or L-cis-diltiazem was quantified by measuring the cGMP-induced current, under steady-state conditions at the end of either −100 mV, −35 mV, or +100 mV pulse, in the presence of diltiazem as required. The amount of diltiazem block (%) is related to the current at the respective cGMP concentration and the amount of current decrease in the presence of diltiazem and was calculated as follow (here exemplified for the 100 μM cGMP-induced current):

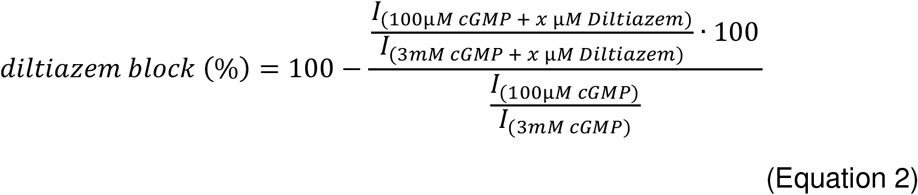

The time courses for channel activation, deactivation (starting after the respective initial delay due to diltiazem removal) and diltiazem block were fitted with a single exponential:

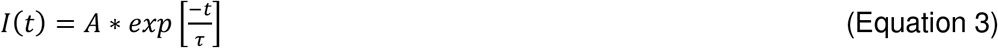

where *A* is the amplitude, *t* the time, and *τ* the time constant for either activation, deactivation, or block.

The time course for diltiazem washout was fitted with a double-exponential function:

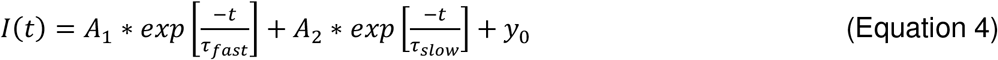

where *A_1_,A_2_* are the amplitudes of the fast and slow components, *t* the time, and τ_fast_ and τ_slow_ the time constants for the fast and slow phase of the diltiazem washout.

For statistical analysis of D- and L-cis-diltiazem effect on retinal CNGCs, we used the two-tailed unpaired Student *t*-test. Figures were prepared using CorelDraw X7 (Corel, Ottawa, Canada).

### Retinal explant culture

To assess the effects of D- and L-cis-diltiazem on calpain activity and photoreceptor degeneration, *rd1* retinas were explanted at post-natal day (P) 5, while retinas from more slowly degenerating *rd10* animals were explanted at P9. The explants were cultured on a polycarbonate membrane (Corning-Costar Transwell permeable support, 24 mm insert, #CLS3412) with complete medium (Gibco R16 medium with supplements) (72). The R16 medium was exchanged every two days with treatment added at either P7 and P9, for *rd1*, or at P11, P13, P15 for *rd10* explants. The cultures were treated with 25, 50, and 100 μM of D- and L-cis-diltiazem, respectively. Cultures were ended on P11 (*rd1*) and P17 (*rd10*) by fixing the cultures with 4% paraformaldehyde (PFA). The explants were embedded in Tissuetek (Sakura Finetek Europe B.V.) and sectioned (12 μm) in a cryostat (ThermoFisher Scientific, CryoStar NX50 OVP, Runcorn UK).

### TUNEL staining

The TUNEL (terminal deoxynucleotidyl transferase dUTP nick end labelling) assay kit (Roche Diagnostics, Mannheim, Germany) was used to label dying cells. Histological sections from retinal explants were dried and stored at −20°C. The sections were rehydrated with phosphate-buffered saline (PBS; 0.1M) and incubated with Proteinase K (1.5 μg/μl) diluted in 50 mM TRIS-buffered saline (TBS; 1 μl enzyme in 7 ml TBS) for 5 mins. This was followed by 3 times 5 minutes TBS washing and incubation with a mixture of 30% HCl and 70% ethanol for 5 min to increase the accessibility of cells to the enzyme. Another 3 times 5 minutes washing was followed by incubation with blocking solution (10% normal goat serum, 1% bovine serum albumin, 1% fish gelatine in phosphate-buffered saline with 0.03% Tween-20). TUNEL staining solution was prepared using 10 parts of blocking solution, 9 parts of TUNEL labelling solution and 1 part of TUNEL enzyme. After blocking, the sections were incubated with TUNEL staining solution overnight at 4° C. Finally, the sections were washed 2 times with PBS, mounted using Vectashield with DAPI (Vector Laboratories Inc, Burlingame, CA, USA) and imaged under a Zeiss (ApoTome.2) microscope for further analysis.

### Calpain-activity assay

This assay allows resolving overall calpain activity *in situ*, on unfixed tissue sections [14]. Retinal tissue sections were incubated and rehydrated for 15 minutes in calpain reaction buffer (CRB) (5.96 g HEPES, 4.85 g KCl, 0.47 g MgCl_2_, 0.22 g CaCl_2_ in 100 ml ddH_2_O; pH 7.2) with 2 mM dithiothreitol (DTT). The tissue sections were incubated for 2 hours at 37°C in CRB with tBOC-Leu-Met-CMAC (5 μM; Thermofisher Scientific, A6520). Afterwards, the tissue sections were washed twice in PBS (5 minutes) and mounted using Vectashield mounting medium (Vector) for immediate visualization under the ZEISS ApoTome2.

### Immunohistochemistry

The cryo-sectioned slides were dried for 30 minutes at 37ºC and hydrated for 15 minutes. The sections were then incubated with blocking solution (10% NGS, 1% BSA and 0.3% PBST) for one hour. The primary antibodies CNGB1 (Sigma-Aldrich, HPA039159; 1:1000), calpain-2 (Abcam, ab39165; 1:300), or caspase-3 (Cell Signalling, 9664; 1:1000) were diluted in blocking solution and incubated overnight at 4ºC. Rinsing with PBS for 3 times 10 minutes each was followed by incubation with secondary antibody (Molecular Probes, AlexaFluor488 (A01134) or AlexaFluor562 (A11036), diluted 1:500 in PBS) for one hour. The sections were further rinsed with PBS for 3 times 10 minutes each and mounted with Vectashield containing DAPI (Vector).

### Microscopy and image analysis in retinal cultures

The images of *ex vivo* retina and organotypic explant cultures were captured using a Zeiss Imager Z.2 fluorescence microscope, equipped with ApoTome2, an Axiocam 506 mono camera, and HXP-120V fluorescent lamp (Carl Zeiss Microscopy, Oberkochen, Germany). The excitation (*λ_Exc_*) / emission (*λ_Em_*.) characteristics of the filter sets used for the different fluorophores were as follows (in nm): DAPI (*λ_Exc_*. = 369 *nm*, *λ_Em_*. = 465 nm), AF488 (*λ_Exc_* = 490 *nm*, *λ_Em_* = 525 nm), and AF562 (*λ_Exc_* = 578 *nm*, *λ_Em_* = 603 *nm*). The Zen 2.3 blue edition software (Zeiss) was used to capture images (both tiled and z-stack, 20x magnification). The data were collected from 7-9 different sections obtained from 3-5 animals. Sections of 12 μm thickness were analysed using 8-12 Apotome Z-planes. The positive cells in the ONL were manually quantified, the ONL area was measured in Zen 2.3 software. The total number of cells in the ONL was calculated using an average cell (nucleus) size and the percent positive cells was determined with respect to the total number of cells in the same ONL area. Values were normalized to control condition (100%).

The relative localization of positive cells within the ONL was assessed by dividing the width of the ONL horizontally into two equal halves (*i.e*. upper and lower half) and manually quantifying the distribution of positive cells in each of the halves. The chance level for cell distribution was 50%. The percent of degenerating photoreceptors localized close to OPL, were analysed by comparing cell count in the lower half of ONL to total positive cells in the ONL.

### Statistical analysis for retinal cultures

Linear mixed-effects models were fitted by restricted maximum likelihood estimation (REML), to assess the significance of the effects in explaining the variations of the dependent variables. Variance inflation factors (VIF) of the predictor variables were calculated and assured to fall well below the common threshold value, indicating no collinearity between them (73). The residuals were confirmed visually to follow a normal distribution, while homoscedasticity (homogeneity of the residual variances) was tested using the Brown-Forsythe test (74) and reported in case of violations.

Figures were prepared using Photoshop CS5 (Adobe, San Jose, CA, USA). Statistical analysis and graph preparation were performed using JMP 15.2.0 (466311, SAS Institute Inc, Cary, NC, USA).

### Two-photon Ca^2+^ imaging

Light stimulus-evoked Ca^2+^ responses were recorded in cone axon terminals using a two-photon (2P) microscope, as previously described (75). In brief, we used adult transgenic HR2.1:TN-XL mice (for details, see above). After dark adaptation for ≥ 1 hour (13), the animal was deeply anesthetized with isoflurane (CP-Pharma, Germany), and then sacrificed by cervical dislocation. All procedures were performed under dim red illumination. Following enucleation of the eyes, the retinas were dissected and vertically sliced (~200 μm) in artificial cerebral spinal fluid (ACSF), which contained (in mM): 125 NaCl, 2.5 KCl, 2 CaCl_2_, 1 MgCl_2_, 1.25 NaH_2_PO_4_, 26 NaHCO_3_, 0.5 L-glutamine, and 20 glucose (Sigma-Aldrich or Merck, Germany) and was maintained at pH 7.4 with carboxygen (95% O_2_, 5% CO_2_). Next, the slices were transferred to the 2P microscope’s recording chamber and superfused with warmed (37°C) ACSF.

The 2P microscope, a customized MOM (Sutter Instruments, Novato, USA; (76), was driven by a mode-locked Ti:Sapphire laser (MaiTai-HP DeepSee; Newport Spectra-Physics, Darmstadt, Germany) tuned to 860 nm. For further technical details on the 2P setup configuration, see (75). TN-XL is a ratiometric FRET-based Ca^2+^ indicator (61), therefore we used two detection channels with the appropriate band-pass (BP) filters (483 BP 32; 535 BP 50; AHF, Tübingen, Germany) to capture both the sensor’s donor (*F_D_*; ECFP) and acceptor fluorescence (*F_A_*; citrine) simultaneously. The relative Ca^2+^ level in the cone terminals was then represented by the ratio *F_A_/F_D_* (*cf*. Fig. 4b, c). Light stimuli were presented using a custom-built stimulator (77) with two band-pass filtered LEDs (UV filter: 360 BP 12; green: 578 BP 10; AHF) mounted below the recording chamber.

Before presenting light flashes and recording cone Ca^2+^ signals, slices were adapted to a constant background illumination equivalent to a photoisomerisation rate of ~10^4^ P*/cone s^-1^ for ≥ 15 seconds. Light stimuli consisted of a series of 1-s bright flashes at 0.25 Hz, evoking similar photoisomerisation rates (~6.5·10^3^ P*s^-1^/cone) in both mouse cone types.

Stock solutions (100mM) of D- and L-cis-diltiazem were prepared in distilled water and stored at 4°C. Prior to each experiment, D- or L-cis-diltiazem dilutions were freshly prepared from the stock in carboxygenated ACSF solution. For bath application, the tissue was perfused with D- or L-cis-diltiazem (25, 50, or 100 μM) added to the bathing solution for ≥ 1 minute before commencing the recording; the perfusion rate was of ~1.5 ml/minute. Drug entry into the recording chamber was confirmed by adding Sulforhodamine 101 (Sigma-Aldrich) to the drug solution.

### Analysis of Ca^2+^-imaging data

To identify the factors (*i.e*. L-cis *vs*. D-cis, concentration) that are significant for predicting the response of a cell during drug treatment (a potential change in AUC), we applied a multivariate linear model (78). The importance of each factor was estimated as its impact on the predictive power of the statistical model. The effect of each factor was considered both individually and in interactions with the other variables, to identify which factor or group of factors is best at modelling the AUC values. The explanatory variables were standardised prior to model fitting, by subtracting the mean and dividing by the standard deviation of the variable. As before, the statistical assumptions of the linear model were evaluated. The VIF for each explanatory variable was found to fall below the common threshold, indicating a lower level of multicollinearity. Visual inspection showed that the residuals were approximately normally distributed. A Brown-Forsythe test indicated that there was heteroscedasticity in the data, though as previously noted these models are robust to such variability. The model also incorporated a random effects term for the recording field, which controlled for recordings where the ROIs were on average higher or lower than the mean across all ROIs in all recordings. Specifically, the modelling showed that (1) more active cells (higher AUC) were more sensitive to the drug application, (2) there was a statistically significant difference between the effects of L- and D-cis-diltiazem on the AUC, and (3) the drug concentration also had a significant effect on AUC.

The effect size is determined using the method for estimating semi-partial R-squared (SPRS) (78) and allowed us to compare the relative impact of each factor in the linear mixed effects model (Table S5). This method also allowed us to evaluate the fit for the whole model (SPRS = 0.368).

## Supplementary Information

The manuscript includes six Supplementary Figures and eight Supplementary Tables.

## ACKNOWLEDGEMENTS

We thank N. Rieger from the Tübingen Institute for Ophthalmic Research as well as K. Schoknecht, S. Bernhardt, A. Kolchmeier from Institute of Physiology II (Jena) for technical assistance. We also thank J. Kusch and K. Benndorf (Institut of Physiology II, Jena) for excellent comments on the manuscript. This work was funded by the ProRetina Foundation and the Deutsche Forschungsgemeinschaft (DFG, German Research Foundation; PA1751/7-1, 8-1 to FPD; EU42/8-1 to TE; TRR 166 ReceptorLight project B01 and Project Number 437036164 to VN).

The authors declare no competing financial interests. FS is General Manager Operations and Head of R & D at Biolog Life Science Institute GmbH & Co. KG. FPD is Chief Scientific Officer for Mireca Medicines GmbH.

## Author contributions

S. Das performed retinal explant cultures, TUNEL and immunostaining, microscopy, analysed data and helped write the manuscript; M. Power performed Ca^2+^-imaging experiments; V. Pop studied the effect of diltiazem on CNGC gating kinetics; K. Groeneveld performed colocalization experiments for heterotetrameric CNGCs; C. Melle performed molecular-biology work; M. Achury performed immunostaining and analysed data; L. Rogerson analysed Ca^2+^-imaging data and performed statistical analysis; T. Strasser performed statistical analysis on immune- and bioassay data; F. Schwede synthesized fluorescent cGMP derivatives; V. Nache performed electrophysiological and optical measurements to study the effect of diltiazem on CNGC; V. Nache, T. Euler, and F. Paquet-Durand designed the experiments, interpreted the data, and prepared the manuscript. All authors edited the manuscript.

## Supplementary Information

**Figure S1:**
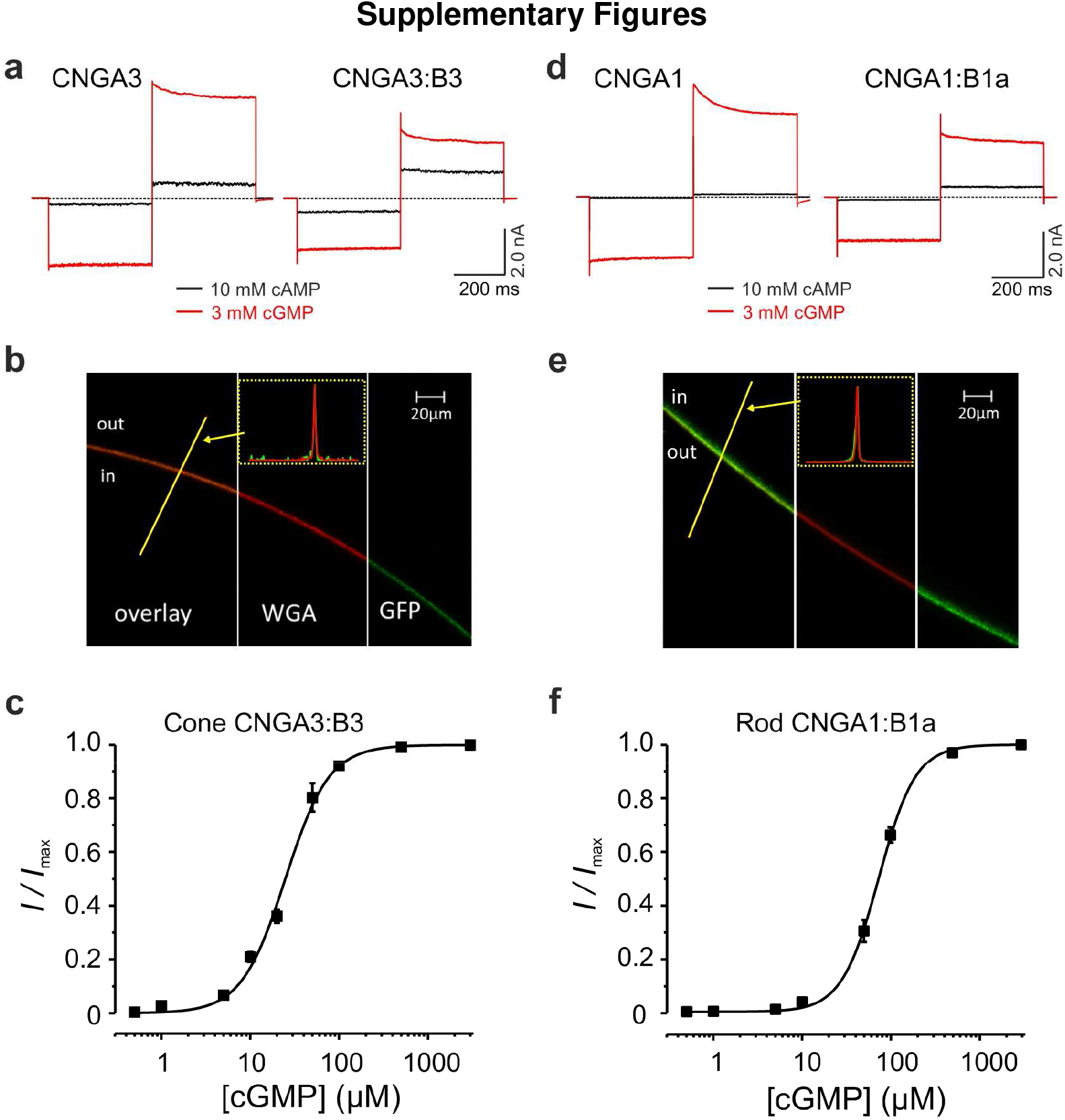
Functional properties of photoreceptor heterotetrameric CNGCs expressed in *Xenopus laevis* oocytes. Representative macroscopic cone (**a**) and rod (**d**) CNGC-current traces from inside-out membrane patches in the presence of 3 mM cGMP (red) and 10 mM cAMP (black). The current traces were elicited by voltage steps from a holding potential of 0 mV to −100, +100 and 0 mV. Leak currents in the absence of cGMP were subtracted for all recordings. For CNGA3 channels the ratio *I*_cAMP_/*I*_cGMP_ was 0.15±0.01 (n=8). CNGB3-subunit incorporation into the CNGA3:B3 channel leads to a significant increase in the cAMP efficacy (*I*_cAMP_/*I*_cGMP_=0.42±0.03, n=6). Similarly, for CNGA1 channels the ratio *I*_cAMP_/*I*_cGMP_ was 0.019±0.005 (n=12), whereas for heterotetrameric CNGA1:B1a channels the ratio was 0.16±0.02 (n=6). (**b, e**) Representative measurements showing confocal images of oocyte membrane expressing heterotetrameric CNGA3:B3-GFP (**b**) and CNGA1:B1a-GFP (**e**) channels (green fluorescence signal). The oocyte plasma membrane was labelled with Alexa Fluor™ 633 WGA (red fluorescence signal). The small insets show fluorescence profiles along the yellow line, perpendicular to the membrane and confirm the colocalization of the labelled channels with the oocyte membrane. For each channel isoform we tested more than 10 oocytes from at least two different oocyte batches. (**c, f**) cGMP-dependent concentration-activation relationships for cone CNGA3:B3 (**c**) and rod CNGA1:B1a (**f**) channels obtained at −35 mV. The currents triggered by subsaturating ligand concentrations were normalized with respect to the maximal current at 3 mM cGMP. The experimental data points, each representing the mean of 5 to 10 measurements, were fitted with Eq. (1) (see also Table S1).

**Figure S2:**
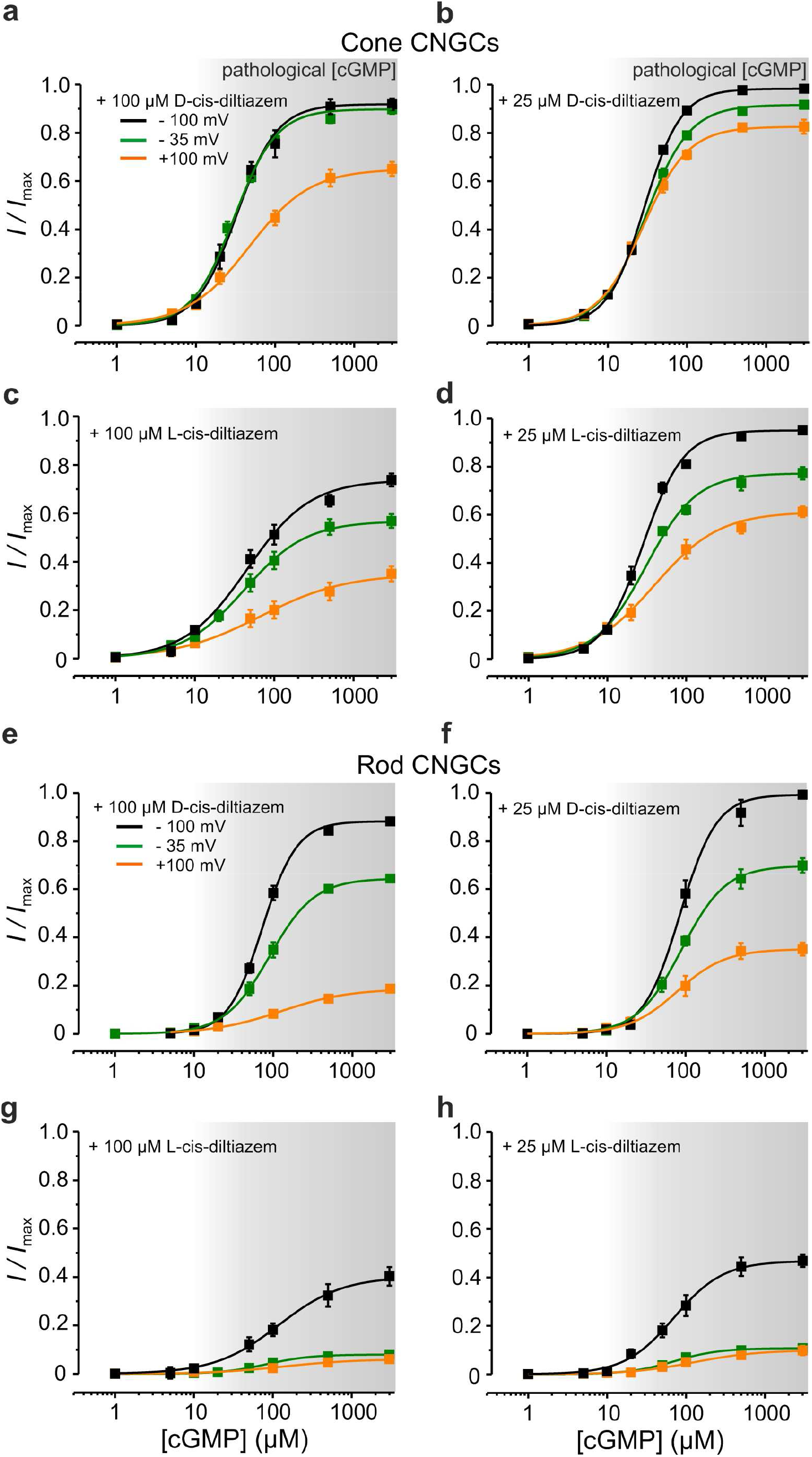
Voltage dependence of D- and L-cis-diltiazem-induced inhibition of photoreceptor CNGCs. cGMP-dependent concentration-activation relationships for cone (**a - d**) and rod (**e - h**) CNGCs in the presence of 100 μM (left) and 25 μM (right) D- and L-cis-diltiazem, respectively, measured at: −100 mV (black symbols), −35 mV (green symbols) and +100 mV (orange symbols). The current amplitudes were normalized with respect to the saturating currents measured in the absence of diltiazem at each individual voltage. The experimental data points were fitted with the Hill equation (Eq. 1). All parameters obtained from the fits are included in Table S1.

**Figure S3:**
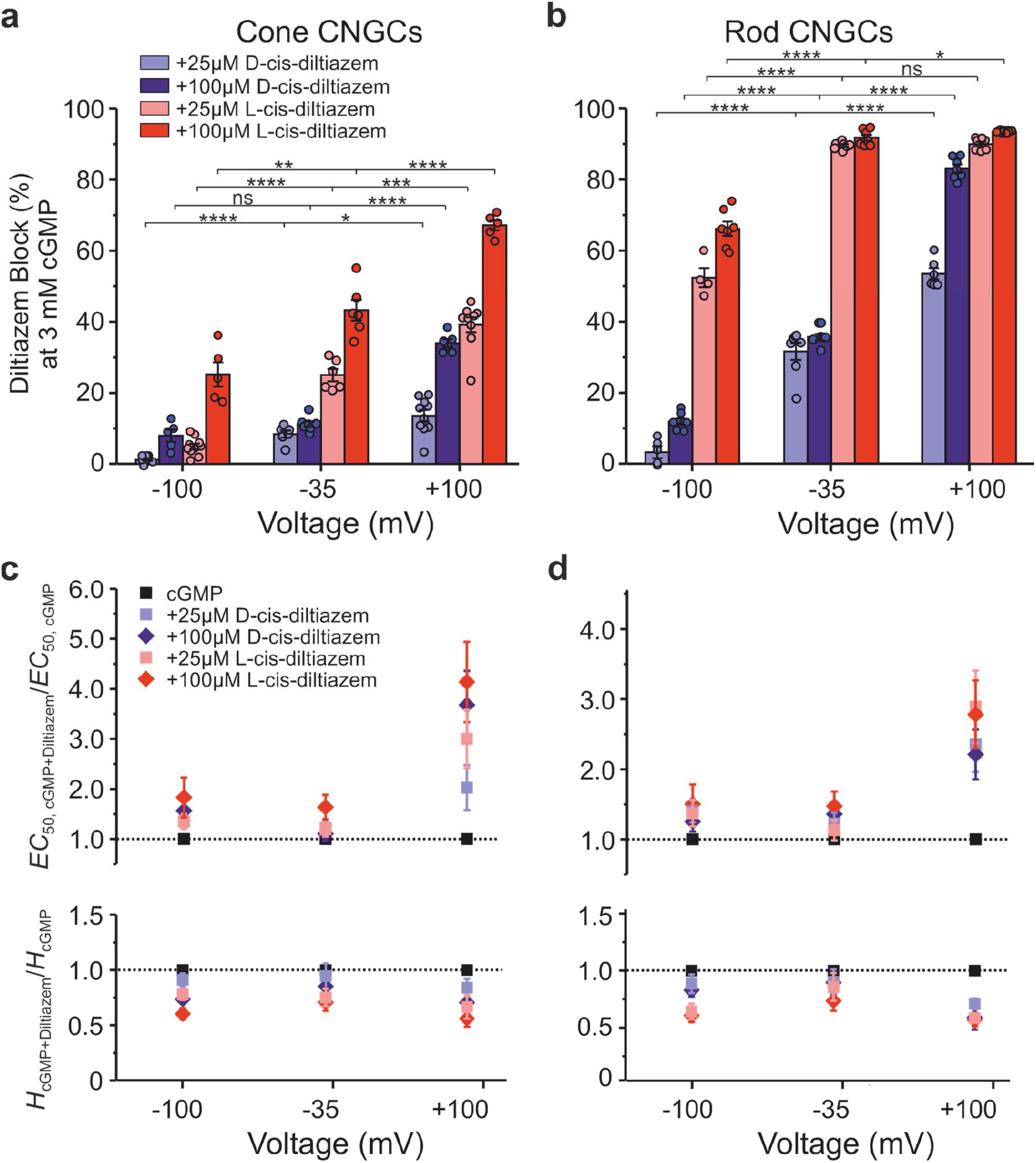
Differential effect of D-cis- and L-cis-diltiazem on CNGC activity and apparent affinity. (**a, b**) D- and L-cis-diltiazem - block of cone and rod CNGC activity triggered by saturating cGMP at three different voltages. The amount of diltiazem block was calculated using Eq. 2. (**c, d**) Effect of D- and L-cis-diltiazem on the channel’s apparent affinity. Shown are the *EC*_50,cGMP+Diltiazem_/*EC*_50,cGMP_- and *H*_cGMP+Diltiazem_ /*H*_cGMP_-ratios in the presence of 25 μM or 100 μM D- or L-cis-Diltiazem at −100 mV, −35 mV and +100 mV. The *EC*_50_- and *H*-values were obtained from the concentration-activation relationships shown in Figs. 1 and S2 (see also Table S1). For statistical analysis see Table S3.

**Figure S4:**
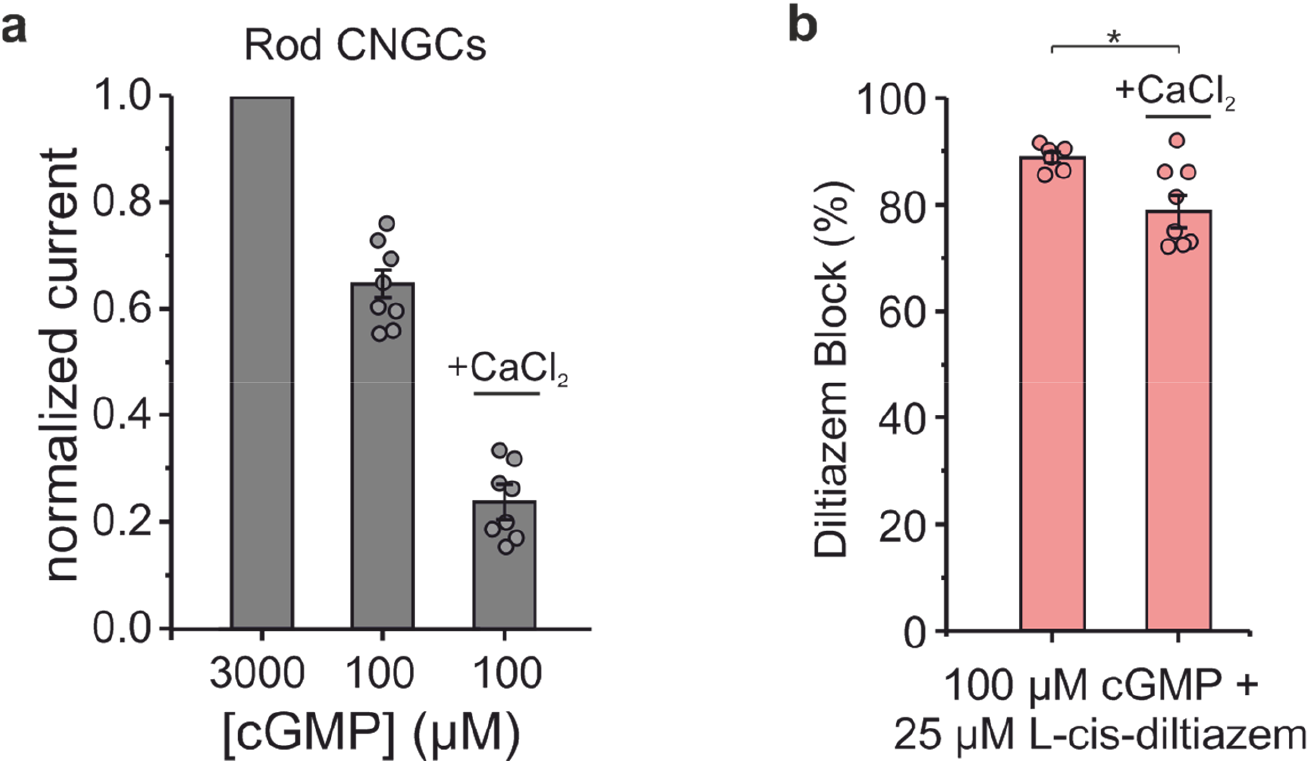
Effect of Ca^2+^ on the blocking effect of L-cis-diltiazem on rod CNGCs. (**a**) The diagram shows normalized rod CNGCs current triggered by 100 μM cGMP, in the absence and in the presence of 1 mM CaCl_2_ in the extracellular solution. The current at 100 μM was normalized with respect to the current in the presence of 3 mM cGMP, under the respective CaCl_2_-conditions (n=9). The channel response to cGMP is much weaker in the presence of Ca^2+^ (*I*_cGMP+caCl2_/*I*_max_ = 0.233±0.03) as it is in its absence (*I*/*I*_max_ = 0.65±0.026). (**b**) L-cis-diltiazem - block of rod CNGC activity triggered by 100 μM cGMP in either the presence or absence of Ca^2+^. The amount of diltiazem block was calculated using Eq. 2. The two-tailed unpaired Student *t*-test was used for the statistical analysis: *p* = 0.034.

**Figure S5:**
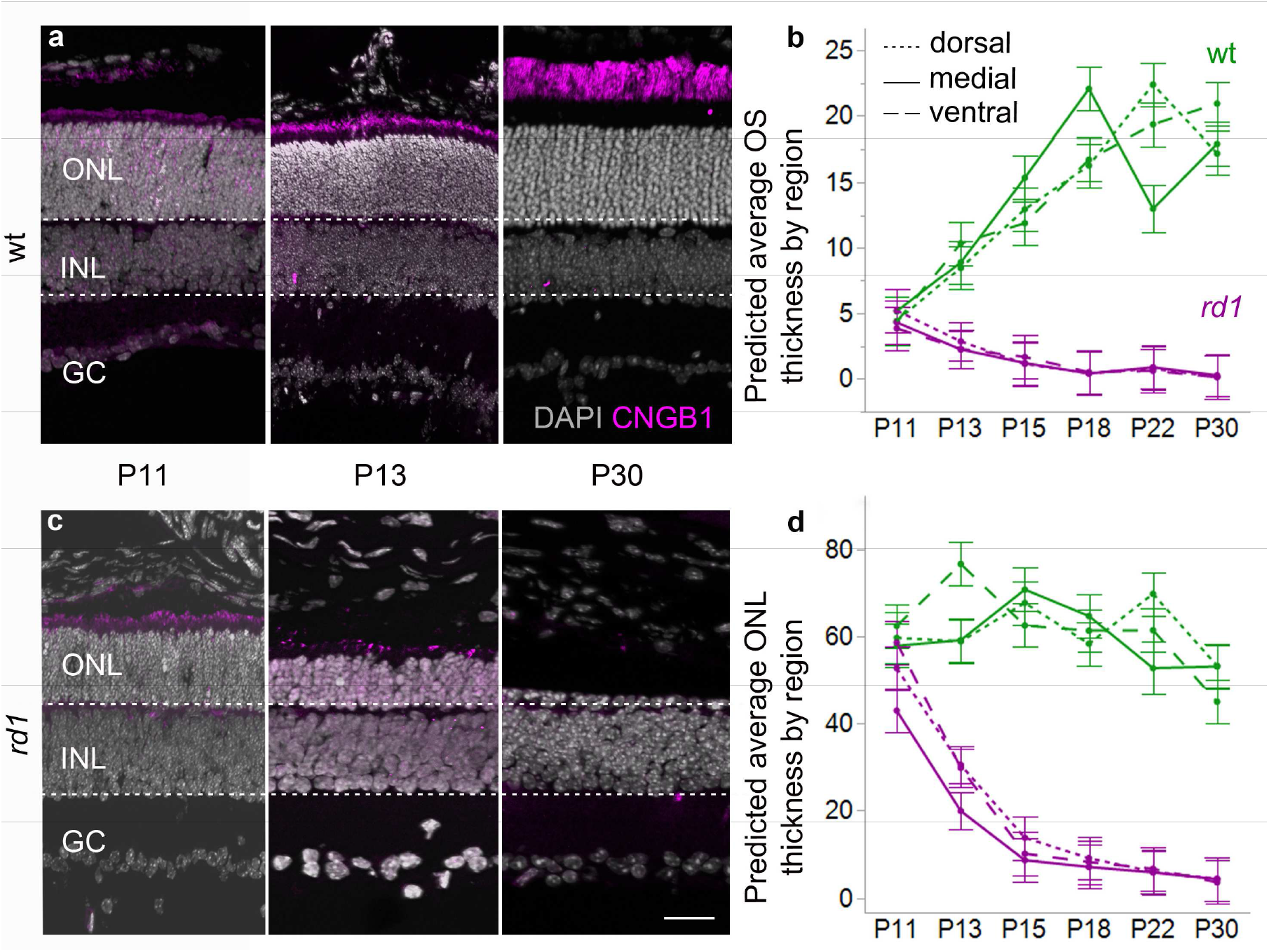
ONL thickness and CNGC expression during *rd1* retinal degeneration. Immunostaining for CNGB1a (magenta) was performed at different post-natal (P) days in wild-type (wt) and *rd1* retina (a,c). The nuclear counterstain (DAPI, grey) indicates outer nuclear layer (ONL), inner nuclear layer (INL), and ganglion cell layer (GC). Dotted, solid and dashed lines in the graph represent dorsal, medial, and ventral mouse retina respectively (b,d). (**a**) In wt retina, CNGB1a immunostaining labelled the photoreceptor outer segments, which grew longer from P11 to P30. (**c**) In *rd1* retina, CNGB1a positive outer segments were visible at P11 and P13 but essentially disappeared by P30. (**d**) The thickness of the ONL in wt retina (green) remained approx. constant between P11 and P30, while *rd1* (magenta) ONL size rapidly diminished after P11. (**b**) Outer segments in wt retina grew longer from P11 to P24 until reaching a plateau at a length of approx. 20 μm. In contrast, *rd1* outer segments, while still comparable to wt at P11, had decreased in length to nearly 0 μm by P24. Images and quantification were obtained from retinal sections from 4-5 different animals per time-point and genotype. Scale bar = 30 μm.

**Figure S6:**
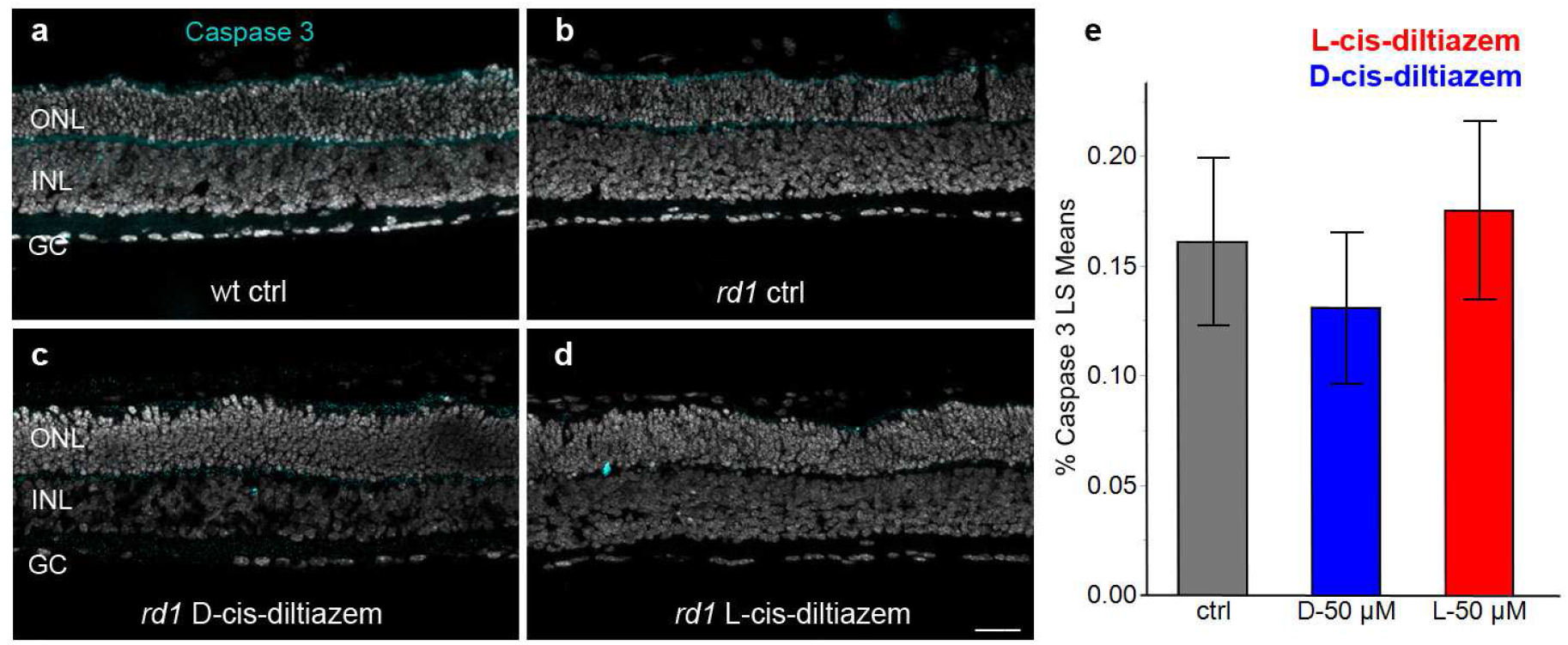
Absence of apoptotic marker during photoreceptor degeneration. Immunostaining for cleaved, activated caspase-3 (turquoise) was performed on *rd1* retinal sections treated with D- and L-cis-diltiazem (50 μM). While caspase-3 immunoreactivity was occasionally found in both outer and inner nuclear layer (ONL, INL), the percentage of caspase-3 positive cells was far lower than the numbers of dying cells (*cf*. Fig. 6). Scale bar = 50 μm.

## Supplementary Tables

**Table S1:**
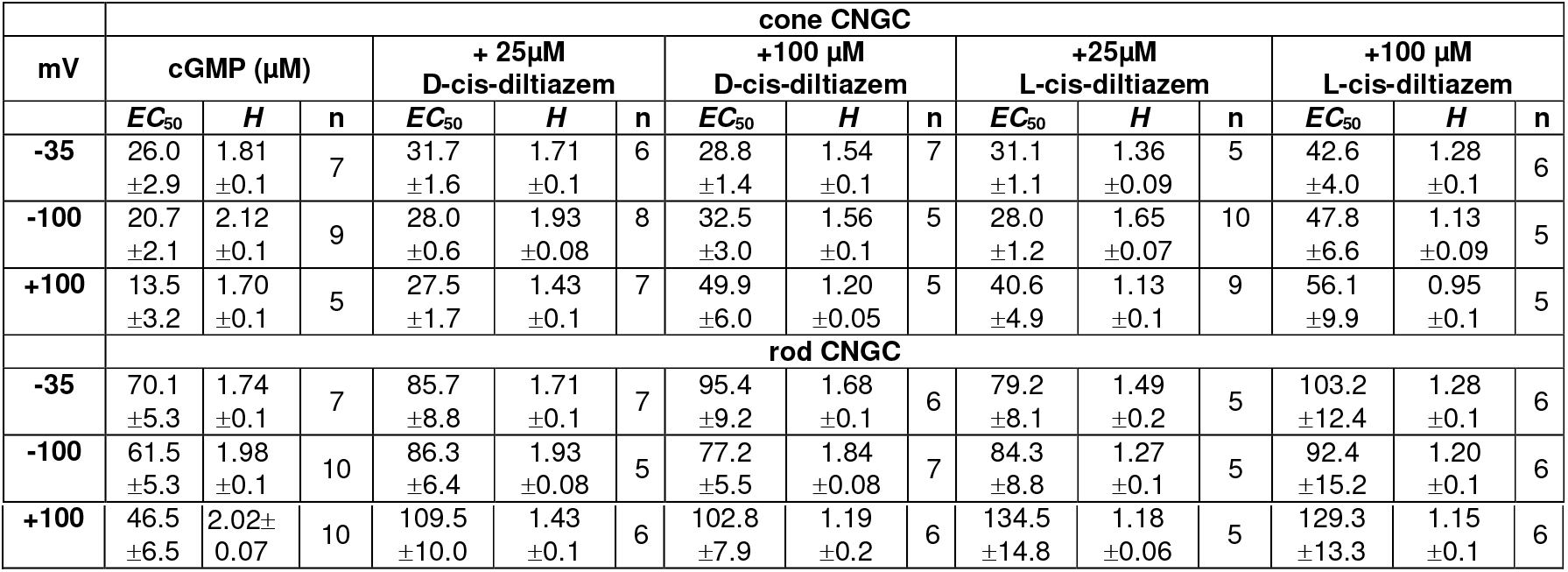
Effect of D- and L-cis-diltiazem on the apparent affinity of rod and cone CNGCs. The *EC*_50_-values and Hill coefficients (*H*, ±SEM) were obtained from the fit of the respective concentrations-activation relationships (n = number of experiments). Two-tailed unpaired Student *t*-test was used to compare the *EC*_50_- and *H*-values in the presence of diltiazem with the ones obtained in its absence.

**Table S2:** Effect of D- and L-cis-diltiazem on the current amplitude of rod and cone CNGCs. The amount of block was determined by comparing the CNGC currents in the presence and in the absence of either D- or L-cis-diltiazem (±SEM, n=5-10) and was calculated using Eq. 2. The comparison between 25 and 100 μm of D- and L-cis-diltiazem, respectively, was performed using the two-tailed unpaired Student’s *t*-test.

| mV | Diltiazem Block (%) of cone CNGC at 3 mM cGMP |  |  |  |  |  |
| --- | --- | --- | --- | --- | --- | --- |
| | + 25 μM<br>D-cis-diltiazem | + 100 μM<br>D-cis-diltiazem | $p$-value | + 25 μM<br>L-cis-diltiazem | + 100 μM<br>L-cis-diltiazem | $p$-value |
| -35 | 8.37 ± 0.97 | 11.2 ± 1.1 | 0.0378 | 25.0 ± 1.6 | 43.2 ± 2.8 | 0.0002 |
| -100 | 1.20 ± 0.3 | 8.02 ± 1.7 | 0.0001 | 4.9 ± 0.8 | 25.3 ± 3.4 | <0.0001 |
| +100 | 13.5 ± 1.6 | 34.0 ± 1.1 | <0.0001 | 39.2 ± 2.2 | 67.2 ± 1.4 | <0.0001 |
| mV | Diltiazem Block (%) of rod CNGC at 3 mM cGMP |  |  |  |  |  |
| | + 25 μM<br>D-cis-diltiazem | + 100 μM<br>D-cis-diltiazem | $p$-value | + 25 μM<br>L-cis-diltiazem | + 100 μM<br>L-cis-diltiazem | $p$-value |
| -35 | 31.5 ± 2.3 | 35.6 ± 0.9 | ns | 89.2 ± 0.37 | 91.5 ± 0.8 | 0.0270 |
| -100 | 3.26 ± 1.7 | 11.9 ± 0.9 | 0.0006 | 52.4 ± 2.7 | 66.2 ± 2.0 | 0.0025 |
| +100 | 53.5 ± 1.6 | 83.1 ± 1.2 | <0.0001 | 90.0 ± 0.65 | 93.5 ± 1.4 | 0.0004 |
| mV | Diltiazem Block (%) of rod CNGC at 100 μM cGMP | | | | | |
| | + 25 μM<br>D-cis-diltiazem | + 100 μM<br>D-cis-diltiazem | $p$-value | + 25 μM<br>L-cis-diltiazem | + 100 μM<br>L-cis-diltiazem | $p$-value |
| -35 | 40.4 ± 3.4 | 46.1 ± 5.3 | ns | 88.4 ± 1.1 | 93.5 ± 1.1 | 0.0021 |

**Table S3:** Statistical analysis of the effect of diltiazem on CNGC *EC*_50_- and *H*-values at different voltages. The respective parameters and number of experiments are listed in Table S1. The *EC*_50_ and *H*-values in the presence of cGMP only were compared with the respective values in presence of cGMP and diltiazem.

| mV | cone CNGC: $p$-value | | | | | | | |
| --- | --- | --- | --- | --- | --- | --- | --- | --- |
| | cGMP + 25μM<br>D-cis-diltiazem | | cGMP + 100 μM<br>D-cis-diltiazem | | cGMP + 25μM<br>L-cis-diltiazem | | cGMP + 100 μM<br>L-cis-diltiazem | |
| | $EC_{50}$ | $H$ | $EC_{50}$ | $H$ | $EC_{50}$ | $H$ | $EC_{50}$ | $H$ |
| -35 | 0.04242 | ns | ns | ns | ns | 0.00677 | 0.00277 | 0.00295 |
| -100 | 0.00042 | ns | 0.000527 | 0.00934 | 0.000291 | 0.00121 | 0.000153 | <0.0001 |
| +100 | 0.000151 | ns | 0.00103 | 0.01494 | 0.00222 | ns | 0.01297 | 0.00636 |
| mV | rod CNGC: $p$-value | | | | | | | |
| | cGMP + 25μM<br>D-cis-diltiazem | | cGMP + 100 μM<br>D-cis-diltiazem | | cGMP + 25μM<br>L-cis-diltiazem | | cGMP + 100 μM<br>L-cis-diltiazem | |
| | $EC_{50}$ | $H$ | $EC_{50}$ | $H$ | $EC_{50}$ | $H$ | $EC_{50}$ | $H$ |
| -35 | ns | ns | 0.02422 | ns | ns | ns | 0.02159 | 0.02111 |
| -100 | 0.00534 | ns | 0.04734 | ns | 0.01729 | 0.00055 | ns | 0.00012 |
| +100 | <0.0001 | <0.0001 | <0.0001 | <0.0001 | <0.0001 | <0.0001 | <0.0001 | <0.0001 |

**Table S4:** Effect of D- and L-cis-diltiazem on the gating kinetics of cone and rod CNGCs. The effect of diltiazem on activation- and deactivation-time constants (τ_act_, τ_deact_ and τ_block_) in the presence of 3 mM cGMP (ms, ±SEM, n=5-9). Two-tailed unpaired Student *t*-test was used for the comparison between time constants obtained in the presence and in the absence of diltiazem.

|  | cone CNGC |  |  |  |  |  |
| --- | --- | --- | --- | --- | --- | --- |
| | $\tau_{act}$ (ms) | *p*-value | $\tau_{deact}$ (ms) | *p*-value | $\tau_{block}$ (ms) | *p*-value |
| cGMP (μM) | 6.8 ± 2.3 | - | 48.2 ± 17.0 | - |  | - |
| + 100 μM<br>D-cis-diltiazem | 7.5 ± 2.6 | ns | 103.6 ± 39.1 | 0.0009 | 154.4 ± 53.4 | ns |
| + 100 μM<br>L-cis-diltiazem | 7.9 ± 2.8 | ns | 94.7 ± 36.8 | 0.0032 | 120.4 ± 38.5 |  |
| rod CNGC |  |  |  |  |  |  |
| cGMP (μM) | 7.6 ± 2.1 | - | 52.1 ± 18.2 | - |  | - |
| + 100 μM<br>D-cis-diltiazem | 9.8 ± 2.6 | ns | 81.2 ± 30.5 | 0.0315 | 139.7 ± 60.2 | ns |
| + 100 μM<br>L-cis-diltiazem | 12.2 ± 4.1 | ns | 123.4 ± 46.1 | 0.0010 | 150.0 ± 56.7 |  |

**Table S5:** Effect of D- and L-cis-diltiazem on light-evoked Ca^2+^ signals in wt cone photoreceptors. The linear modelling identified the variables that significantly predict the data. The area-under-the-curve (AUC) in the control condition was significant and had the largest effect size, with semi-partial R-squared (SPRS) equal to 0.195 (*p* < 0.0001). The drug treatment and the drug concentration were both significant. There was also a statistically significant interaction between the AUC in the control condition, the drug treatment, and the drug concentration. Since their confidence intervals overlap, we cannot state which of these model components had the greatest effect size. There was neither a significant interaction between the AUC in the control condition and the drug treatment, nor between the AUC in the control condition and the drug concentration. (*cf*. Fig. 4).

| Component | *p*-value | Effect-Size | ES-Lower-CI | ES-Upper-CI |
| --- | --- | --- | --- | --- |
| Model (all components) |  | 0.368 | 0.319 | 0.422 |
| AUC (control) | <0.0001 | 0.195 | 0.147 | 0.247 |
| Treatment (drug) | 0.00123 | 0.05 | 0.023 | 0.085 |
| Treatment (concentration) | 0.3416 | 0.039 | 0.016 | 0.072 |
| AUC (control) x treatment (drug) | 0.119 | 0.003 | 0 | 0.017 |
| AUC (control) x treatment (conc.) | 0.171 | 0.002 | 0 | 0.015 |
| AUC (control) x treatment (drug) x treatment (conc.) | <0.0001 | 0.021 | 0.005 | 0.047 |

**Table S6:** Analysis of the variability of OS length and ONL thickness in *rd1* and wt. Results of the linear mixed-effects models with the dependent variables OS length and ONL thickness. The models’ residuals followed a normal distribution, while the Brown-Forsythe test indicated a violation of the assumption of homoscedasticity for both models. However, linear mixed-effects models estimates have been shown to be robust against such violations (79).

|  | OS length<br>n = 109, R<sup>2</sup><sub>adj.</sub> = 0.96 |  | ONL thickness<br>n = 109, R<sup>2</sup><sub>adj.</sub> = 0.93 |  |
| --- | --- | --- | --- | --- |
| Fixed effect | F-statistic | p-value | F-statistic | p-value |
| genotype | F(1, 25.25) = 0.0078 | 0.9304 | F(1, 25.99) = 2.6450 | 0.1159 |
| Time-point | F(5, 24.7) = 4.4213 | 0.0052 | F(5, 24.77) = 15.1946 | < 0.0001 |
| Retinal position | F(2, 48.36) = 0.2982 | 0.7435 | F(2, 49.39) = 2.9380 | 0.0623 |
| genotype x time-point | F(5, 24.7) = 13.2699 | < 0.0001 | F(5, 24.77) = 12.0885 | < 0.0001 |
| genotype x retinal position | F(2, 48.36) = 0.3245 | 0.7245 | F(2, 49.39) = 0.9156 | 0.4070 |
| timepoint x retinal position | F(10, 47.94) = 3.8401 | 0.0007 | F(10, 48.41) = 2.0258 | 0.0508 |
| genotype x time-point x retinal position | F(10, 47.94) = 4.2248 | 0.0003 | F(10, 48.41) = 1.3344 | 0.2397 |

**Table S7:** Analysis of cell death markers using linear mixed-effects models. Shown are the effects that explain the variability of the dependent variables TUNEL, calpain activity, calpain-2 positive cells, as well as localization of TUNEL positive cells within the ONL. All models included the animal as a random effect to account for repeated measures. Numbers in brackets indicate the total number of animals used per genotype, n represents the number of observations used in the model. Normality of residuals was assessed visually; heterogeneity of residual variances (homoscedasticity) was tested with the Brown-Forsythe test. Linear mixed-effects models have been shown to be robust against violations of model assumptions.

| Dependent variable | Genotype | Fixed effect | Normality of residuals | Homo-scedasticity | F-statistic | p-value |
| --- | --- | --- | --- | --- | --- | --- |
| TUNEL | wt (35)<br>R<sup>2</sup><sub>adj.</sub> = .80<br>n = 336 | Concentration<sup>1</sup> | Yes | No | F(3, 17.92) = 20.7656 | <0.0001 |
|  |  | Treatment |  |  | F(1, 303.74) = 0.171 | 0.6795 |
|  |  | Concentration x Treatment |  |  | F(3, 22.8) = 29.6038 | <0.0001 |
|  | *rd1* (36)<br>R<sup>2</sup><sub>adj.</sub> = .88<br>n = 331 | Concentration<sup>1</sup> | Yes | No | F(3, 24.82) = 37.8570 | < 0.0001 |
|  |  | Treatment |  |  | F(1, 306.63) = 0.0787 | 0.7792 |
|  |  | Concentration x Treatment |  |  | F(3, 27.75) = 31.0649 | <0.0001 |
|  | *rd10* (10)<br>R<sup>2</sup><sub>adj.</sub> = .84<br>n = 112 | Concentration<sup>4</sup> | Yes | No | F(1, 8.11) = 25.9134 | <0.0009 |
|  |  | Treatment |  |  | F(1, 100.96) = 0.0026 | 0.9598 |
|  |  | Concentration x Treatment |  |  | F(1, 10.43) = 23.5461 | <0.0006 |
| Calpain activity | wt (11) & *rd1* (11)<br>R<sup>2</sup><sub>adj.</sub> = .86<br>n = 143 | Concentration<sup>4</sup> | Yes | No | F(1, 16.41) = 99.2752 | <0.0001 |
|  |  | Treatment |  |  | F(2, 16.41) = 0.793 | 0.4691 |
|  |  | Concentration x Treatment |  |  | F(2, 16.41) = 2.055 | 0.1598 |
| Calpain-2 | wt (9) & *rd1* (9)<br>R<sup>2</sup><sub>adj.</sub> = .83<br>n = 117 | Genotype | Yes | No | F(1, 12.14) = 9.0927 | 0.0106 |
|  |  | Treatment |  |  | F(2, 12.14) = 20.2775 | 0.0001 |
|  |  | Genotype x Treatment |  |  | F(2, 12.14) = 2.2535 | 0.1471 |
| Caspase-3 | *rd1* (11)<br>R<sup>2</sup><sub>adj.</sub> = .04<br>n = 58 | Treatment | Yes | Yes | F(2, 7.15) = 0.3799 | 0.6970 |
| ONL localisation TUNEL | *rd1* (9)<br>R<sup>2</sup><sub>adj.</sub> = .72<br>n = 53 | Treatment | Yes | No | F(2, 10.11) = 49.4033 | <0.0001 |

Treatment: {D-cis-diltiazem, L-cis-diltiazem}, <sup>1</sup>{0, 25, 50, 100 µM}, <sup>2</sup>{0, 25 µM}, <sup>3</sup>{0, 50 µM}, <sup>4</sup>{0, 100 µM}

**Table S8:** Post-hoc analysis of the linear mixed-effects models. Results of contrast tests comparing the least-square means, which resulted from the linear mixed-effects models shown in Table S7.

| | Contrast<br>LS means<br>[95% confidence interval] (%) | | LS means<br>diff. ± SE (%) | F-statistic | p-value |
| --- | --- | --- | --- | --- | --- |
| **ONL**<br>**localisation**<br>**TUNEL** | *rd1* ctrl<br>59.98<br>[54.04, 65.92] | *rd1* D-25 μM<br>57.44<br>[52.05, 62.83] | 2.54 ± 3.79 | F(1, 16.33) = 0.4511 | 0.5112 |
| | | *rd1* L-25 μM<br>85.03<br>[79.70, 90.36] | 25.05 ± 3.40 | F(1, 10.42) = 54.2025 | < 0.0001 |
| **Calpain-2** | *rd1* ctrl<br>1.82<br>[1.20, 2.44] | *rd1* D-50 μM<br>0.68<br>[0.07, 1.30] | 1.13 ± 0.40 | F(1, 12.52) = 7.9008 | 0.0152 |
| | | *rd1* L-50 μM<br>2.73<br>[2.11, 3.35] | 0.91 ± 0.40 | F(1, 12.69) = 5.0979 | 0.0423 |
| | wt ctrl<br>0.52<br>[0.09, 1.13] | wt L-50 μM<br>2.04<br>[1.43, 2.66] | 1.52 ± 0.39 | F(1, 11.87) = 14.7372 | 0.0024 |
| **TUNEL** | *rd1* ctrl<br>98.10<br>[50.88, 145.32] | *rd1* D-100 μM<br>180.85<br>[111.06, 250.63] | 82.75 ± 41.14 | F(1, 28.11) = 4.0454 | 0.0540 |
| | *rd1* ctrl<br>101.96<br>[54.54, 149.38] | *rd1* L-100 μM<br>661.96<br>[593.66, 730.26] | 560.00 ± 40.99 | F(1, 26.68) = 191.1994 | <0.0001 |
| | *rd10* ctrl<br>90.68<br>[201.44, 382.81] | *rd10* D-100 μM<br>401.33<br>[111.58, 691.07] | 310.60 ± 178.75 | F(1, 8.10) = 3.0200 | 0.1200 |
| | *rd10* ctrl<br>93.60<br>[198.61, 385.80] | *rd10* L-100 μM<br>1403.14<br>[1060.80, 1745.48] | 1310.00 ± 199.71 | F(1, 9.25) = 42.9966 | <0.0001 |
| | wt ctrl<br>105.35<br>[58.38, 152.32] | wt D-100 μM<br>112.44<br>[43.49, 181.39] | 7.10 ± 39.87 | F(1, 19.17) = 0.0316 | 0.8607 |
| | wt ctrl<br>98.28<br>[51.36, 145.21] | wt L-50 μM<br>458.14<br>[406.36, 509.93] | 359.90 ± 33.31 | F(1, 18.59) = 116.6931 | <0.0001 |
| | | wt L-100 μM<br>420.42<br>[366.06, 474.78] | 322.10 ± 34.59 | F(1, 21.63) = 86.7207 | <0.0001 |

## REFERENCES

1. Kennan A, Aherne A, Humphries P. Light in retinitis pigmentosa. Trends Genet 21, 103–110 (2005)

2. Narayan DS, Wood JP, Chidlow G, Casson RJ. A review of the mechanisms of cone degeneration in retinitis pigmentosa. Acta Ophthalmol 94, 748–754 (2016)

3. Hamel CP. Cone rod dystrophies. Orphanet J Rare Dis 2, 7 (2007)

4. Das S, Chen Y, Yan J, Christensen G, Belhadj S, Tolone A, et al. The role of cGMP-signalling and calcium-signalling in photoreceptor cell death: perspectives for therapy development. Pflugers Arch (2021)

5. Barabas P, Cutler PC, Krizaj D. Do calcium channel blockers rescue dying photoreceptors in the Pde6b (rd1) mouse? Adv Exp Med Biol 664, 491–499 (2010)

6. Wetzel RK, Arystarkhova E, Sweadner KJ. Cellular and Subcellular Specification of Na,K-ATPase α and β Isoforms in the Postnatal Development of Mouse Retina. J Neurosci 19, 9878–9889 (1999)

7. Waldner DM, Bech-Hansen NT, Stell WK. Channeling Vision: Ca(V)1.4-A Critical Link in Retinal Signal Transmission. BioMed Res Int 7, 1–14 (2018)

8. Ingram NT, Sampath AP, Fain GL. Membrane conductances of mouse cone photoreceptors. J Gen Physiol 152, (2020)

9. Paquet-Durand F, Beck S, Michalakis S, Goldmann T, Huber G, Muhlfriedel R, et al. A key role for cyclic nucleotide gated (CNG) channels in cGMP-related retinitis pigmentosa. Hum Mol Genet 20, 941–947 (2011)

10. Fox DA, Poblenz AT, He LH. Calcium overload triggers rod photoreceptor apoptotic cell death in chemical-induced and inherited retinal degenerations. Ann Ny Acad Sci 893, 282–285 (1999)

11. Vallazza-Deschamps G, Cia D, Gong J, Jellali A, Duboc A, Forster V, et al. Excessive activation of cyclic nucleotide-gated channels contributes to neuronal degeneration of photoreceptors. Eur J Neurosci 22, 1013–1022 (2005)

12. Bowes C, Li T, Frankel WN, Danciger M, Coffin JM, Applebury ML, et al. Localization of a retroviral element within the rd gene coding for the beta subunit of cGMP phosphodiesterase. PNAS 90, 2955–2959 (1993)

13. Wei T, Schubert T, Paquet-Durand F, Tanimoto N, Chang L, Koeppen K, et al. Light-driven calcium signals in mouse cone photoreceptors. J Neurosci 32, 6981–6994 (2012)

14. Kulkarni M, Trifunovic D, Schubert T, Euler T, Paquet-Durand F. Calcium dynamics change in degenerating cone photoreceptors. Hum Mol Genet 25, 3729–3740 (2016)

15. Schon C, Paquet-Durand F, Michalakis S. Cav1.4 L-Type Calcium Channels Contribute to Calpain Activation in Degenerating Photoreceptors of rd1 Mice. PLoS One 11, e0156974 (2016)

16. Sothilingam V, Garcia-Garrido M, Jiao K, Buena-Atienza E, Sahaboglu A, Trifunovic D, et al. Retinitis Pigmentosa: Impact of different Pde6a point mutations on the disease phenotype. Hum Mol Genet 24, 5486–5499 (2015)

17. Hart J, Wilkinson MF, Kelly ME, Barnes S. Inhibitory action of diltiazem on voltage-gated calcium channels in cone photoreceptors. Exp Eye Res 76, 597–604 (2003)

18. Stern JH, Kaupp UB, MacLeish PR. Control of the light-regulated current in rod photoreceptors by cyclic GMP, calcium, and l-cis-diltiazem. PNAS 83, 1163–1167 (1986)

19. Frasson M, Sahel JA, Fabre M, Simonutti M, Dreyfus H, Picaud S. Retinitis pigmentosa: rod photoreceptor rescue by a calcium-channel blocker in the rd mouse. Nat Med 5, 1183–1187 (1999).

20. Fox DA, Poblenz AT, He L, Harris JB, Medrano CJ. Pharmacological strategies to block rod photoreceptor apoptosis caused by calcium overload: a mechanistic target-site approach to neuroprotection. Eur J Ophthalmol 13, 44–56 (2003)

21. Pawlyk BS, Sandberg MA, Berson EL. Effects of IBMX on the rod ERG of the isolated perfused cat eye: antagonism with light, calcium or L-cis-diltiazem. Vision Res 31, 1093–1097 (1991)

22. Pearce-Kelling SE, Aleman TS, Nickle A, Laties AM, Aguirre GD, Jacobson SG, et al. Calcium channel blocker D-cis-diltiazem does not slow retinal degeneration in the PDE6B mutant rcd1 canine model of retinitis pigmentosa. Mol Vis 7, 42–47 (2001)

23. Pawlyk BS, Li T, Scimeca MS, Sandberg MA, Berson EL. Absence of photoreceptor rescue with D-cis-diltiazem in the rd mouse. Invest Ophthalmol Vis Sci 43, 1912–1915 (2002)

24. Shuart NG, Haitin Y, Camp SS, Black KD, Zagotta WN. Molecular mechanism for 3:1 subunit stoichiometry of rod cyclic nucleotide-gated ion channels. Nat Commun 2, 457 (2011)

25. Peng C, Rich ED, Varnum MD. Subunit configuration of heteromeric cone cyclic nucleotide-gated channels. Neuron 42, 401–410 (2004)

26. Cote RH, Brunnock MA. Intracellular cGMP concentration in rod photoreceptors is regulated by binding to high and moderate affinity cGMP binding sites. J Biol Chem 268, 17190–17198 (1993)

27. Nakatani K, Yau KW. Calcium and light adaptation in retinal rods and cones. Nature 334, 69–71 (1988)

28. Frings S, Seifert R, Godde M, Kaupp UB. Profoundly different calcium permeation and blockage determine the specific function of distinct cyclic nucleotide-gated channels. Neuron 15, 169–179 (1995)

29. Picones A, Korenbrot JI. Permeability and interaction of Ca2+ with cGMP-gated ion channels differ in retinal rod and cone photoreceptors. Biophys J 69, 120–127 (1995)

30. Nache V, Eick T, Schulz E, Schmauder R, Benndorf K. Hysteresis of ligand binding in CNGA2 ion channels. Nat Commun 4, 2866 (2013)

31. Biskup C, Kusch J, Schulz E, Nache V, Schwede F, Lehmann F, et al. Relating ligand binding to activation gating in CNGA2 channels. Nature 446, 440–443 (2007)

32. Nache V, Wongsamitkul N, Kusch J, Zimmer T, Schwede F, Benndorf K. Deciphering the function of the CNGB1b subunit in olfactory CNG channels. Sci Rep 6, 29378 (2016)

33. Arango-Gonzalez B, Trifunović D, Sahaboglu A, Kranz K, Michalakis S, Farinelli P, et al. Identification of a common non-apoptotic cell death mechanism in hereditary retinal degeneration. PLoS One 9, e112142 (2014)

34. Powers TA, Rogin C. MarkeTrak 10: History and Methodology. Semin Hear 41, 3–5 (2020)

35. Nicholson DW, Ali A, Thornberry NA, Vaillancourt JP, Ding CK, Gallant M, et al. Identification and inhibition of the ICE/CED-3 protease necessary for mammalian apoptosis. Nature 376, 37–43 (1995)

36. Vighi E, Trifunovic D, Veiga-Crespo P, Rentsch A, Hoffmann D, Sahaboglu A, et al. Combination of cGMP analogue and drug delivery system provides functional protection in hereditary retinal degeneration. PNAS 115, E2997–E3006 (2018)

37. Bocchero U, Tam BM, Chiu CN, Torre V, Moritz OL. Electrophysiological Changes During Early Steps of Retinitis Pigmentosa. Invest Ophthalmol Vis Sci 60, 933–943 (2019)

38. Haynes LW. Block of the cyclic GMP-gated channel of vertebrate rod and cone photoreceptors by l-cis-diltiazem. J Gen Physiol 100, 783–801 (1992)

39. McLatchie LM, Matthews HR. Voltage-dependent block by L-cis-diltiazem of the cyclic GMP-activated conductance of salamander rods. Proc Biol Sci 247, 113–119 (1992)

40. Tang L, Gamal El-Din TM, Lenaeus MJ, Zheng N, Catterall W. Structural Basis for Diltiazem Block of a Voltage-gated Ca2+ Channel. Mol Pharmacol 96, 485–492 (2019)

41. Zhao Y, Huang G, Wu J, Wu Q, Gao S, Yan Z, et al. Molecular Basis for Ligand Modulation of a Mammalian Voltage-Gated Ca(2+) Channel. Cell 177, 1495–1506 (2019)

42. Zheng X, Fu Z, Su D, Zhang Y, Li M, Pan Y, et al. Mechanism of ligand activation of a eukaryotic cyclic nucleotide-gated channel. Nat Struct Mol Biol 27, 625–634 (2020)

43. Xue J, Han Y, Zeng W, Wang Y, Jiang Y. Structural mechanisms of gating and selectivity of human rod CNGA1 channel. Neuron 109, 1302–1313 (2021)

44. Fain GL, Lisman JE. Light, Ca2+, and photoreceptor death: new evidence for the equivalent-light hypothesis from arrestin knockout mice. Invest Ophthalmol Vis Sci 40, 2770–2772 (1999)

45. Olshevskaya EV, Ermilov AN, Dizhoor AM. Factors that affect regulation of cGMP synthesis in vertebrate photoreceptors and their genetic link to human retinal degeneration. Mol Cell Biochem 230, 139–147 (2002)

46. Johnson JE, Jr., Perkins GA, Giddabasappa A, Chaney S, Xiao W, White AD, et al. Spatiotemporal regulation of ATP and Ca2+ dynamics in vertebrate rod and cone ribbon synapses. Mol Vis 13, 887–919 (2007)

47. Ames A, III. Energy requirements of CNS cells as related to their function and to their vulnerability to ischemia: a commentary based on studies on retina. Can J Physiol Pharmacol 70, S158–S64 (1992)

48. Wegierski T, Kuznicki J. Neuronal calcium signaling via store-operated channels in health and disease. Cell Calcium 74, 102–111 (2018)

49. Johnson MT, Gudlur A, Zhang X, Xin P, Emrich SM, Yoast RE, et al. L-type Ca2+; channel blockers promote vascular remodeling through activation of STIM proteins. PNAS 117, 17369 (2020)

50. Saraiva N, Prole DL, Carrara G, Johnson BF, Taylor CW, Parsons M, et al. hGAAP promotes cell adhesion and migration via the stimulation of store-operated Ca2+ entry and calpain 2. J Cell Biol 202, 699–713 (2013)

51. Olshevskaya EV, Ermilov AN, Dizhoor AM. Factors that affect regulation of cGMP synthesis in vertebrate photoreceptors and their genetic link to human retinal degeneration. Mol Cell Biochem 230, 139–147 (2002)

52. Paquet-Durand F, Hauck SM, van Veen T, Ueffing M, Ekström P. PKG activity causes photoreceptor cell death in two retinitis pigmentosa models. J Neurochem 108, 796–810 (2009)

53. Power M, Das S, Schutze K, Marigo V, Ekstrom P, Paquet-Durand F. Cellular mechanisms of hereditary photoreceptor degeneration - Focus on cGMP. Prog Retin Eye Res 74, 100772 (2020)

54. Krizaj D, Copenhagen DR. Compartmentalization of calcium extrusion mechanisms in the outer and inner segments of photoreceptors. Neuron 21, 249–256 (1998)

55. Spencer M, Detwiler PB, Bunt-Milam AH. Distribution of membrane proteins in mechanically dissociated retinal rods. Invest Ophthalmol Vis Sci 29, 1012–1020 (1988)

56. Koch S, Sothilingam V, Garcia Garrido M, Tanimoto N, Becirovic E, Koch F, et al. Gene therapy restores vision and delays degeneration in the CNGB1(-/-) mouse model of retinitis pigmentosa. Hum Mol Genet 21, 4486–4496 (2012)

57. Bareil C, Hamel CP, Delague V, Arnaud B, Demaille J, Claustres M. Segregation of a mutation in CNGB1 encoding the beta-subunit of the rod cGMP-gated channel in a family with autosomal recessive retinitis pigmentosa. Hum Genet 108, 328–334 (2001)

58. Wissinger B, Gamer D, Jägle H, Giorda R, Marx T, Mayer S, et al. CNGA3 mutations in hereditary cone photoreceptor disorders. Am J Hum Genet 69, 722–737 (2001)

59. Wutz K, Sauer C, Zrenner E, Lorenz B, Alitalo T, Broghammer M, et al. Thirty distinct CACNA1F mutations in 33 families with incomplete type of XLCSNB and Cacna1f expression profiling in mouse retina. Eur J Hum Genet 10, 449–456 (2002)

60. Sanyal S, Bal AK. Comparative light and electron microscopic study of retinal histogenesis in normal and rd mutant mice. Z Anat Entwicklungsgesch 142, 219–238 (1973)

61. Mank M, Reiff DF, Heim N, Friedrich MW, Borst A, Griesbeck O. A FRET-Based Calcium Biosensor with Fast Signal Kinetics and High Fluorescence Change. Biophys J 90, 1790–1796 (2006)

62. Kaupp UB, Niidome T, Tanabe T, Terada S, Bonigk W, Stuhmer W, et al. Primary structure and functional expression from complementary DNA of the rod photoreceptor cyclic GMP-gated channel. Nature 342, 762–766 (1989)

63. Korschen HG, Illing M, Seifert R, Sesti F, Williams A, Gotzes S, et al. A 240 kDa protein represents the complete beta subunit of the cyclic nucleotide-gated channel from rod photoreceptor. Neuron 15, 627–636 (1995)

64. Yu WP, Grunwald ME, Yau KW. Molecular cloning, functional expression and chromosomal localization of a human homolog of the cyclic nucleotide-gated ion channel of retinal cone photoreceptors. FEBS Lett 393, 211–215 (1996)

65. Peng C, Rich ED, Thor CA, Varnum MD. Functionally important calmodulin-binding sites in both NH2- and COOH-terminal regions of the cone photoreceptor cyclic nucleotide-gated channel CNGB3 subunit. J Biol Chem 278, 24617–24623 (2003)

66. Liman ER, Tytgat J, Hess P. Subunit stoichiometry of a mammalian K+ channel determined by construction of multimeric cDNAs. Neuron 9, 861–871 (1992)

67. Shammat IM, Gordon SE. Stoichiometry and arrangement of subunits in rod cyclic nucleotide-gated channels. Neuron 23, 809–819 (1999)

68. Jonas P. High-speed solution switching using piezo-based micropositioning stages. In: Sakmann B, Neher E (eds) Single-channel recording. 2nd edn. (Springer US, Plenum Press, New York 1995) pp xxii–700

69. Thon S, Schulz E, Kusch J, Benndorf K. Conformational Flip of Nonactivated HCN2 Channel Subunits Evoked by Cyclic Nucleotides. Biophys J 109, 2268–2276 (2015)

70. Zheng J, Zagotta WN. Patch-clamp fluorometry recording of conformational rearrangements of ion channels. Sci STKE 2003, PL7 (2003)

71. Nache V, Zimmer T, Wongsamitkul N, Schmauder R, Kusch J, Reinhardt L, et al. Differential regulation by cyclic nucleotides of the CNGA4 and CNGB1b subunits in olfactory cyclic nucleotide-gated channels. Sci Signal 5, ra48 (2012)

72. Belhadj S, Tolone A, Christensen G, Das S, Chen Y, Paquet-Durand F. Long-Term, Serum-Free Cultivation of Organotypic Mouse Retina Explants with Intact Retinal Pigment Epithelium. J Vis Exp, e61868 (2020)

73. Hair JF, Anderson RE, Tatham RL, Black W. Multivariate data analysis New York. NY: Macmillan. (1995)

74. Nobre JS, da Motta Singer J. Residual analysis for linear mixed models. Biom J 49, 863–875 (2007)

75. Kulkarni M, Schubert T, Baden T, Wissinger B, Euler T, Paquet-Durand F. Imaging Ca2+ dynamics in cone photoreceptor axon terminals of the mouse retina. J Vis Exp, e52588 (2015)

76. Euler T, Hausselt SE, Margolis DJ, Breuninger T, Castell X, Detwiler PB, et al. Eyecup scope--optical recordings of light stimulus-evoked fluorescence signals in the retina. Pflugers Arch 457, 393–414 (2009)

77. Baden T, Schubert T, Chang L, Wei T, Zaichuk M, Wissinger B, et al. A Tale of Two Retinal Domains: Near-Optimal Sampling of Achromatic Contrasts in Natural Scenes through Asymmetric Photoreceptor Distribution. Neuron 80, 1206–1217 (2013)

78. Nakagawa S, Schielzeth H. A general and simple method for obtaining R2 from generalized linear mixed-effects models. Methods Ecol Evol 4, 133–142 (2013)

79. Schielzeth H, Dingemanse NJ, Nakagawa S, Westneat DF, Allegue H, Teplitsky C, et al. Robustness of linear mixed-effects models to violations of distributional assumptions. Methods Ecol Evol 11, 1141–1152 (2020)

